# Vaccine induction of CD4-mimicking broadly neutralizing antibody precursors in macaques

**DOI:** 10.1101/2023.03.05.531154

**Authors:** Kevin O. Saunders, James Counts, Victoria Stalls, Robert Edwards, Kartik Manne, Xiaozhi Lu, Bhishem Thakur, Katayoun Mansouri, Yue Chen, Rob Parks, Maggie Barr, Laura Sutherland, Joena Bal, Nicholas Havill, Haiyan Chen, Emily Machiele, Nolan Jamieson, Bhavna Hora, Megan Kopp, Katarzyna Janowska, Kara Anasti, Chuancang Jiang, Sravani Venkatayogi, Amanda Eaton, Rory Henderson, Christopher Barbosa, S. Munir Alam, Sampa Santra, Drew Weissman, M. Anthony Moody, Derek W. Cain, Ying Tam, Mark Lewis, Wilton B. Williams, Kevin Wiehe, David Montefiori, Priyamvada Acharya, Barton F. Haynes

## Abstract

The CD4 binding site (CD4bs) is a conserved epitope on HIV-1 envelope (Env) that can be targeted by protective broadly neutralizing antibodies (bnAbs). HIV-1 vaccines have not elicited CD4bs bnAbs for many reasons, including the CD4bs is occluded by glycans, immunogen expansion of appropriate naïve B cells, and selection of functional antibody mutations. Here, we demonstrate immunization of macaques with a CD4bs-targeting immunogen elicits neutralizing bnAb precursors with structural and genetic features of CD4-mimicking bnAbs. Structures of the CD4bs nAbs bound to HIV-1 Env demonstrated binding angles similar to human bnAbs and heavy chain second complementarity determining region-dependent binding characteristic of all known human CD4-mimicking bnAbs. Macaque nAbs were derived from variable and joining gene segments orthologous to the genes of human V_H_1-46-class bnAbs. This vaccine study initiated the B cells from which derive CD4bs bnAbs in primates, accomplishing the key first step in development of an effective HIV-1 vaccine.

## INTRODUCTION

Broadly neutralizing antibodies (bnAbs) can protect against sensitive viruses in humans and animal models of HIV-1 infection ^1–4^, and are a primary goal of HIV-1 vaccine development^5^. BnAbs target one of seven conserved epitopes on HIV-1 Env^6^. Among these Env conserved sites is the binding site for CD4 (CD4bs)^6^. Monoclonal antibody isolation from people living with HIV-1 has identified two classes of bnAbs that mimic CD4 in the manner in which they bind Env^6–10^. The first class of bnAbs are derived from the VH1-2*02 germline gene segment and includes VRC01, CH31, and 3BNC117 ^10, 11^. This type of CD4 mimicking antibody uses beta strands in its heavy chain second complementarity determining region (HCDR2) to recapitulate the beta strands of CD4 allowing both proteins to fit similarly into the CD4bs^7,10^. Extensive studies have been conducted to characterize the frequency of this type of CD4 mimicking antibody in the human repertoire, and found the precursors to be relatively rare^12^. The second class of CD4 mimicking bnAbs are derived from the VH1-46 gene segment and include bnAbs such as CH235.12, 8ANC131, and 1-18^6,11,13,14^. Like VH1-2*02-derived bnAbs, these antibodies mimic CD4 using beta strands in their HCDR2^13,15^. Vaccine design efforts have been focused on the VRC01 class antibodies ^16–20^, and relatively less investigation has aimed to elicit VH1-46 class bnAbs^15,21,22^. However, the potent and broad neutralization of VH1-46-derived bnAbs, the lack of insertions and deletions in their genes, heterogenous light chain gene usage, and normal LCDR3 lengths make this type of bnAb a desirable vaccine design target.

It has been proposed that the first step in eliciting bnAbs with vaccination is for Env immunogens to bind the bnAb precursor—termed the unmutated common ancestor (UCA) antibody of the bnAb B cell clone^23–25^. Therefore, one approach to eliciting VH1-46-class CD4bs bnAbs is to engineer high affinity Env immunogens that bind to the VH1-46-derived, unmutated B cell receptor of naïve B cells that target the CD4bs ^5,23–26^. To enable design of such immunogens for VH1-46-class CD4bs bnAbs, we previously isolated the CH235 bnAb lineage and contemporaneous HIV-1 Env sequences from the same individual, named CH505^13,27,28^. We engineered an Env that bound the UCA antibody of the CH235 lineage (CH235 UCA) by introducing N279K and G458Y substitutions into the HIV-1 Env inferred to have initiated the infection in the CH505 individual ^15,21^. This engineered Env, called CH505 M5.G458Y, induced serum autologous neutralizing CD4bs antibodies and selected for functional somatic mutations needed for neutralization breadth in CH235 UCA knock-in mice^21^. Similarly, immunization of rhesus macaques with M5.G458Y Env trimer conjugated to ferritin nanoparticles generated CD4bs serum neutralizing antibodies^21^. These macaque serum CD4bs antibodies showed hallmarks of the CH235 lineage in that the neutralizing antibodies were dependent upon the N279K and/or G458Y substitutions engineered into the Env to promote CH235 UCA binding ^21^. However, it was unknown whether these serum neutralizing CD4bs antibodies were IGHV1-derived, CD4 mimicking neutralizing antibodies.

Here, we vaccinated rhesus macaques with M5.G458Y Env trimers and a novel lipid nanoparticle adjuvant and elicited serum CD4bs autologous tier 2 virus neutralizing antibodies. The serum antibodies bound to HIV-1Env with orientations comparable to CH235. Isolated monoclonal neutralizing CD4bs antibodies from multiple macaques utilized rhesus VH gene segments orthologous to human VH1-46 and had angles of approach to Env similar to the CH235 bnAb. A high-resolution structure of one of these vaccine-induced antibodies showed it mimicked CD4 utilizing structural features similar to VH1-46 bnAbs. The hallmark amino acids identified to mediate binding of human CD4bs bnAbs were also functionally required in rhesus macaque CD4bs nAbs. Lastly, the inferred precursor of this vaccine-induced CD4bs nAb bound to the vaccine immunogen consistent with the vaccine goal of targeting specific VH1-46-like germline antibodies. Thus, this study demonstrates induction of CD4 mimicking, CD4bs nAbs in rhesus macaques with paratope structures, immunogenetics, and neutralization signatures recapitulating human VH1-46-type bnAb precursors.

## RESULTS

### Vaccine induction of serum CD4bs nAbs

To elicit VH1-46 bnAb-like antibody responses, three rhesus macaques were vaccinated six times with the CH505 M5.G458Y HIV-1 Env engineered to bind to the CH235 UCA^15,21^ (**Figure 1A**). M5.G458Y Env was enriched for Man_5_GlcNAc_2_ glycans and adjuvanted with a lipid nanoparticle that has been shown to boost antibody production for both mRNA and protein immunogens^29^. Serum IgG responses to the autologous or vaccine-matched Env arose after a single immunization and peaked after two immunizations, withMan_5_GlcNAc_2_-enrichment on Env improving serum IgG binding (**Figure 1B**). We performed competition assays with serum and CH235.12 to determine whether the Env-specific serum antibody response was targeted to the CD4bs. Serum antibody throughout the first five immunizations showed increasing ability to block the binding of CD4bs bnAb CH235.12, but exhibited low blocking of N332-glycan bnAb 2G12 (**Figure 1C**). Thus, immunization elicited a substantial CD4bs binding antibody response compared to N332 glycan-directed antibodies.

**Figure 1.**
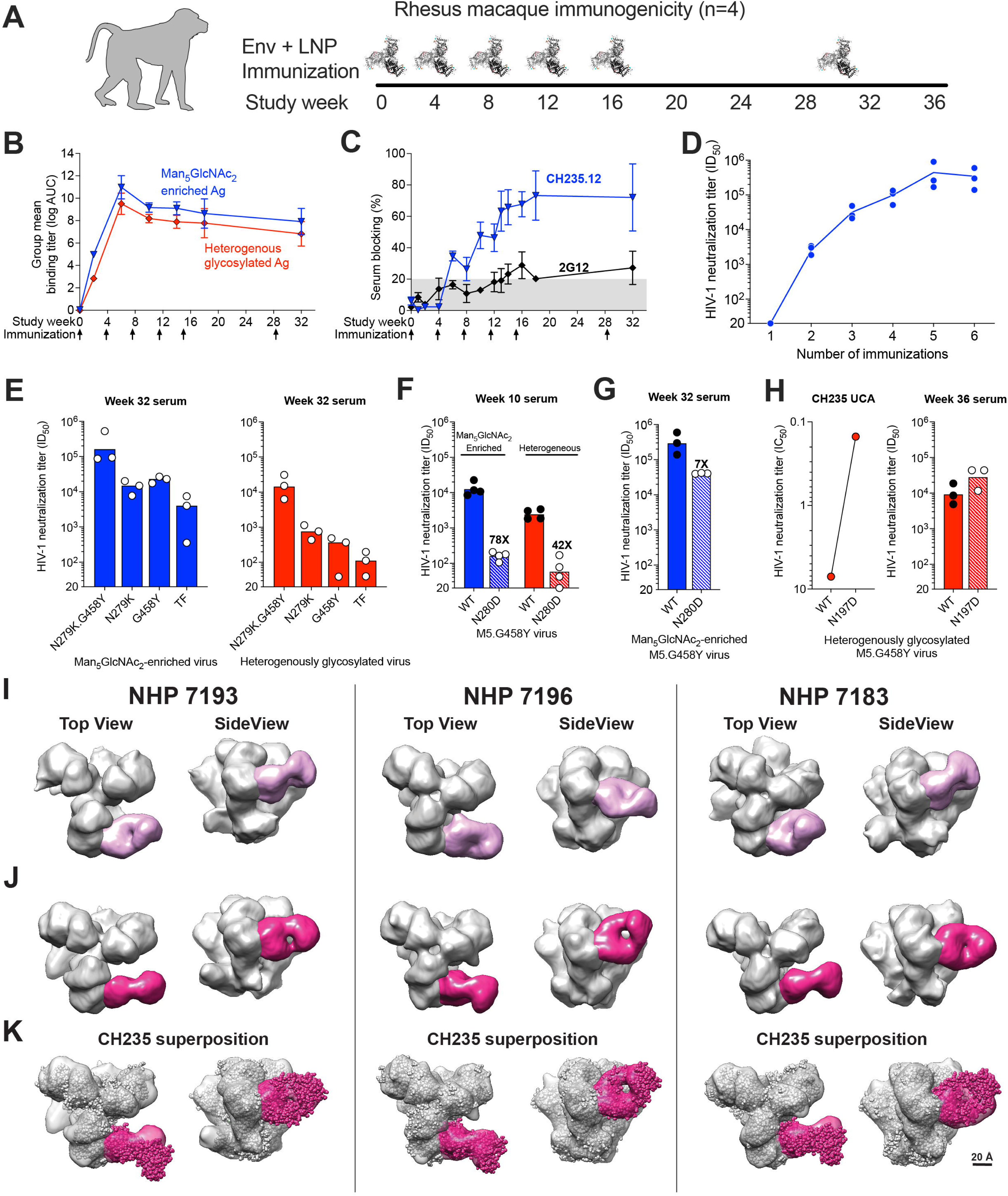
CH505.M5.G458Y stabilized gp140 Trimers Induce Serum CD4 Binding Site-Directed Antibodies In Rhesus Macaques. **(A)** Vaccination of rhesus macaques with Man_5_GlcNAc_2_-enriched CH505.M5.G458Y Env trimers formulated with ionizable lipid nanoparticles. **(B)** Serum IgG binding magnitude to Man_5_GlcNAc_2_-enriched (blue) and heterogeneously glycosylated (red) vaccine-matched Env. Values are reported as the group mean and standard error for the area under the log_10_ transformed concentration curve (logAUC) over time. Immunization time points indicated by arrows. **(C)** Serum blocking of CD4bs bnAb CH235.12 (blue) and N332 glycan bnAb 2G12 (gray) binding to CH505.M5.G458Y Env. **(D)** Serum neutralization ID50 titer against vaccine-matched tier 2 pseudotyped virus increases over the course of vaccination. Trend line shows group geometric mean (n=3 macaques). **(E)** Week 32 (post-6 immunizations) neutralization activity depends upon germline-targeting mutations N279K (M5) and G458Y, as well as Man_5_GlcNAc_2_ enrichment. In **E-H**, different glycoforms of pseudovirus are color-coded as in **B**. Bars represent group geometric mean titers. Values for individual macaques are shown as symbols. **(F,G)** Weeks 10 (F, post 3 immunizations) and 32 (G) serum neutralization of CH505.M5.G458Y pseudovirus is greatly diminished in the presence of CD4bs KO mutation N280D. Values reported above the bars in F and G are fold change in the group geometric mean titer. **(H)** Removal of the N-linked glycosylation site at Env position 197, which shields the CD4bs, improves serum neutralization of heterogeneously glycosylated CH505.M5.G458Y pseudovirus by both the CH235 UCA and vaccinated macaque serum. **(I-K)** 3D reconstruction of negative stain electron microscopy images of serum-derived Fabs bound to CH505.M5.G458Y Env trimers. (**I**)Top and side view of serum CD4bs antibodies with binding orientations such that both Fab chains are visible when looking down from the trimer. (**J**) Top and side view of serum CD4bs antibodies with binding orientations similar to CH235. The Fab orientation is rotated 90 degrees compared to the Fabs in **I**. **(K)** Superposition of the CH235-bound Env structure (spheres representation) into the observed density for serum Fabs bound to Env (surface representation).

The neutralizing antibody ID50 titer against the vaccine-matched virus was high, peaking at a group geometric mean of 1:339,853 serum dilution after the fifth immunization (**Figure 1D**). Heterologous neutralization was modest against CH235-sensitive viruses (**Figure S1A,B**). CH235 lineage antibodies at the beginning of affinity maturation exhibit a neutralization signature where they potently neutralize viruses that have both N279K (also called M5) and G458Y substitutions, moderately neutralize viruses with either N279K or G458Y, and weakly neutralize viruses lacking both substitutions^15^. After six immunizations serum neutralizing antibodies were most potent when virion-associated Env included both N279K and G458Y. Neutralization was weaker if only one these substitutions was present, and weakest if neither substitution was present (**Figure 1E**). Thus, the serum neutralization signature matched that of the CH235 lineage^15,21^. The serum neutralization of M5.G458Y virus was also sensitive to a CD4bs amino acid change from Asn to Asp at position 280 (N280D), which is known to knockout CH235 lineage antibody neutralization early in development^21^. As the CH235 lineage evolves it becomes less affected by N280D substitution (**Figure S1C**). Serum neutralization showed the same pattern, since after three immunizations the N280D substitution reduced neutralization titer 78-fold, but after six immunizations only reduced neutralization titer 7-fold (**Figure 1F and G**).

Glycans proximal to the CD4bs can hinder neutralization by CD4bs antibodies^30,31^. The CH235 lineage antibodies bind in the vicinity of the N197 glycan. While the N197 glycan is disordered in the cryo-EM structure of the CH235 UCA complex (PDB ID 6UDA)^15^, cryo-EM structures of CH235.12 bound to CH505.N279K (or M5) SOSIP Env trimers, showed well-defined density for glycan N197 making close contact with a region of the antibody heavy chain near K19T, E81D and T70Y substitutions (**Figure S2 and Table S1**), suggesting accommodation of glycan N197 by the maturing CH235 lineage through the acquisition of somatic mutations. A K19T substitution in CH235 UCA improved its binding to CH505 Envs, and removal of the N197 glycosylation site improved CH235 UCA antibody neutralization (**Figure S2 and S3**)^15^. Consistent with elicitation of CH235-like antibodies, vaccinated macaque serum neutralization potency also increased upon removal of the N197 glycosylation site (**Figure 1H**).

We next visualized Env epitopes targeted by the elicited antibodies using antigen binding fragments (Fabs) prepared from total IgG purified from the serum of each vaccinated macaque, and performed negative stain electron microscopic polyclonal epitope mapping (EMPEM) of the Fabs bound to the M5.G458Y Env trimer. Three dimensional (3D) classifications from all three animals showed robust polyclonal Fab binding to the CD4bs, as well as binding to the trimer base, the N611 glycan epitope, and the V1/V3 epitope (**Figure S4**). Masked 3D classifications focused on the CD4bs demonstrated all NHPs developed antibodies that bind to the CD4bs with a horizontal approach such that the flat plane of the Fab is approximately parallel to the plane of the page when seen in top view (**Figure 1I**). All NHPs also developed antibodies that bound the CD4bs with an angle of approach more similar to canonical VH1-46-type of CD4bs bnAbs (**Figure 1J**). Rigid-body fitting of the Env-CH235 UCA complex into the EMPEM maps shows that this latter subset of serum CD4bs antibodies have binding modes similar to CH235 UCA at this resolution (**Figure 1K**). Hence, structural studies indicated vaccine elicitation of CD4bs antibody responses targeting the CH235 epitope.

### Induction of lymph node TFH responses

In addition to the serum antibody response, we analyzed vaccine-induced TFH cell responses in lymph nodes harvested prior to vaccination and one week after the 3rd and 4th immunizations (weeks 9 and 13 after immunization). We quantified Env-specific TFH cells using an Activation-Induced Marker Assay^32,33^, using an overlapping peptide pool spanning CH505TF gp140 to stimulate lymph node cells ex vivo (**Supplementary Figure 5A**). Env-specific TFH cells were not detected in lymph node biopsies from any monkey at pre-immunization time points but were readily detected in all four NHP after the 3rd and 4th immunizations (**Figure S5B**), indicative of vaccine-induced Env-specific CD4 T cell help for humoral responses. Similarly, Env-specific B cells were undetectable in lymph nodes prior to vaccination, but steadily increased in frequency after the 3rd and 4th immunizations (**Figure S5C and S5D**). Vaccination with CH505 M5 G458Y GnTI-SOSIP nanoparticles/LNP elicited robust humoral responses that were supported by cognate CD4 T cell help.

### Distinct phenotypic classes of recombinant CD4bs neutralizing antibodies

To compare vaccine-induced antibodies more specifically to VH1-46 bnAbs, we isolated antibody sequences from blood memory B cells for phenotypic and genotypic characterization as recombinant IgG antibodies (Abs) (**Figure 2A and S6A**). M5.G458Y Env trimer-specific single B cells were sorted from two rhesus macaques (RM7193 and RM7196), and their corresponding B cell receptors were sequenced. Approximately 30% of the immunoglobulin heavy chain variable regions were derived from IGHV1 in each macaque (**Figure S6B**). Using the KIMDB database of *Maccaca mulatta* sequences we assigned IGHV gene segments to immunoglobulin sequences. Three clonal lineages were derived from rhesus IGHV1-105, which is orthologous to human IGHV1-46 (**Figure 2B**). The most abundant macaque IGHV used by the antigen-specific B cells was IGHV1-84, which was only 78% identical to both rhesus IGHV1-105 and human IGHV1-46 (**Figure 2B and S6C**). The three VH1-105 utilizing antibodies displayed common distributions of mutation percentages and CDR3 lengths **(Figure S6D and S6E)**.

**Figure 2.**
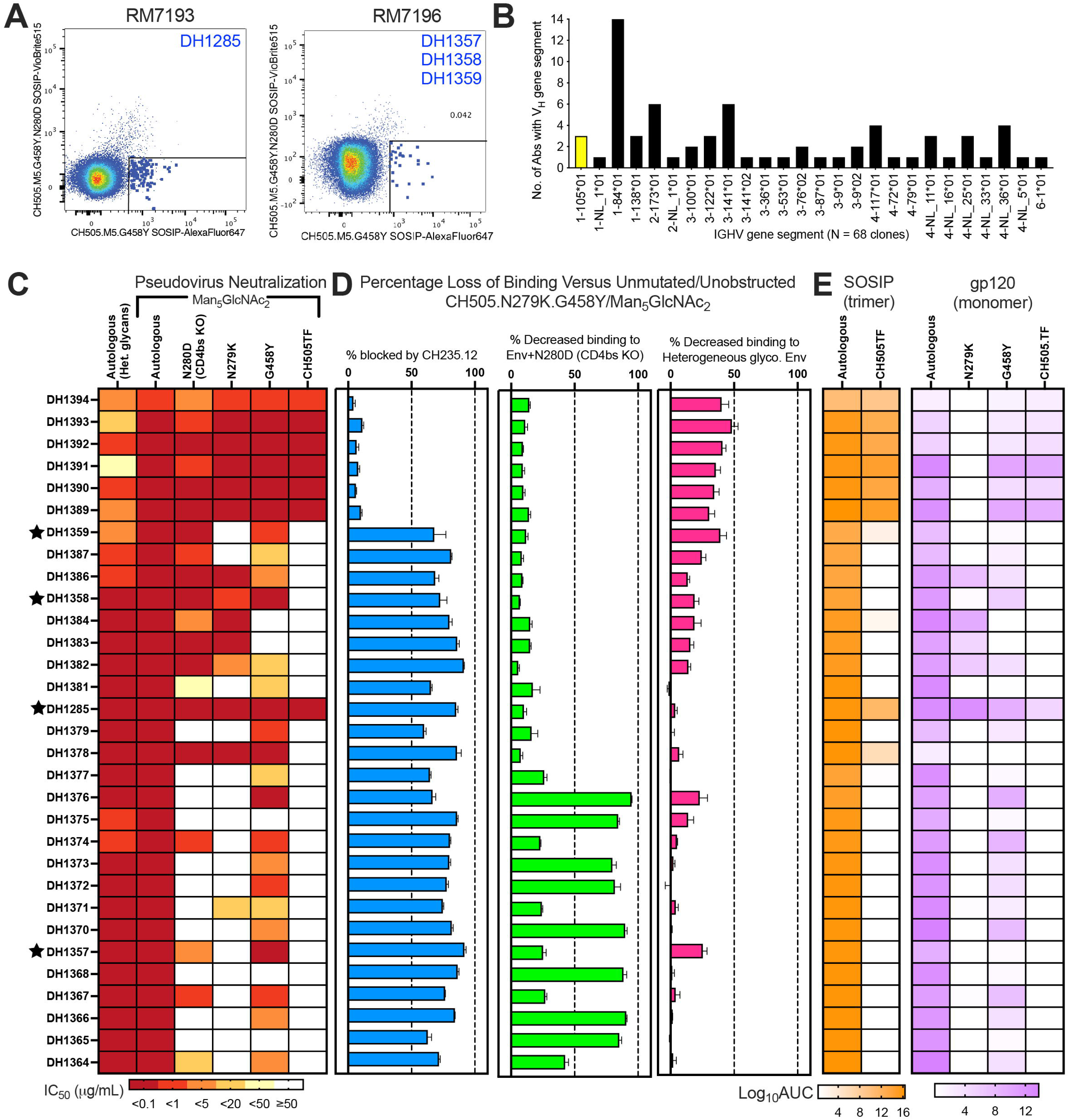
CH505.M5.G458Y-Specific Monoclonal Antibodies from Vaccinated Macaques Demonstrate CD4bs-Directed Binding and Neutralization. **(A)** Fluorescence-activated sorting of Env reactive single B cells that lack binding in the presence of CD4 KO substitution N280D. **(B)** IGHV gene segments used by unique B cell clones isolated from NHP7193 and 7196. IGHV1-105 is highlighted yellow since it is orthologous to the human IGHV1-46 gene segment used by CD4bs bnAbs. **(C)** Pseudotyped virus neutralization IC50 titer against autologous virus with and without Man_5_GlcNAc_2_ enrichment. Neutralization was mapped to the CD4bs by removal of CD4bs bnAb targeting substitutions (N279K or G458Y) or CD4bs knockout substitution (N280D). **(D)** Percent loss of macaque antibody binding due to competition with CH235.12 binding to Env (blue), the presence of CD4bs knockout substitution N280D (green), or heterogenous Env glycosylation (pink) as determined by ELISA. Blocking magnitude or decrease in binding is determined relative to binding to CH505.M5.G458Y/GnT1^-^. Error bars represent the standard deviation of three replicates. **(E)** Binding reactivity, reported as logAUC, for SOSIP trimers (orange) (N=3 independent experiments) or gp120 versions of HIV-1 Env by ELISA (N=2 independent experiments). Neutralizing antibodies that are derived from IGHV1-105 and compete with CH235.12 for binding to envelope are marked with stars beside their names.

Fifty-three recombinant antibodies were selected for further study based on derivation from IGHV1 gene segments and/or initial Env binding screens. Antibodies were assessed to determine whether Env reactivity was dependent on Man_5_GlcNAc_2_ glycosylation or amino acid substitutions at N280, N279K, or G458Y. Of the fifty-three antibodies, thirty-one (58%) neutralized the autologous, vaccine-matched virus, Man_5_GlcNAc_2_-enriched M5.G458Y (**Figure 2C and S7**). Overall, Man_5_GlcNAc_2_ enrichment enhanced antibody neutralization, but had a minor effect on binding to soluble Env trimers (**Figure 2C and 2D**). Twenty-one of thirty-one (68%) neutralizing antibodies exhibited at least a five-fold reduction in neutralization in the presence of a N280D substitution (**Figure 2C**). The same percentage of antibodies were dependent on N279K or G458Y for neutralization of the vaccine-matched virus. Generally, both of these phenotypes were concordant with observed loss of binding to N280D versions of M5.G458Y soluble Env or to Env gp120 without N279K or G458Y (**Figure 2D**). Twenty-four antibodies showed a 50% or greater reduction in Env binding magnitude in the presence of CH235.12 (**Figure 2D**). Altogether, the binding phenotype coupled with the competition with CH235.12 for binding to Env indicated the majority (68%) of nAbs exhibited a CD4bs specificity similar to CH235 lineage antibodies (**Figure 2C-2E**).

Seven of the fifty-three antibodies (DH1285 and DH1389-DH1394) were capable of neutralizing all of the CH505 viruses tested regardless of the presence of N279K or G458Y(**Figure 2C and Figure S7A**). For these seven nAbs, Man_5_GlcNAc_2_ enrichment improved M5.G458Y recombinant Env binding as well as pseudovirus neutralization (**Figure 2C and 2D**). Additionally, these seven recombinant antibodies bound to both M5.G458Y and CH505 TF soluble Env trimers **(Figure 2E)**. However, only 1 of the 7 nAbs, DH1285, competed with CH235.12 for binding to Env and bound to Env gp120 monomers with or without N279K and G458Y substitutions **(Figure 2D)**. These results suggested DH1285 was a CD4bs antibodies that no longer required N279K or G458Y, whereas the other 6 Man_5_GlcNAc_2_-enriched CH505 TF nAbs targeted a different epitope (**Figure 2 and S7**).

### IGHV1-105-derived macaque nAbs exhibit a CH235-like Env binding mode

From the 53 monoclonal antibodies, we found four nAbs (DH1285, DH1357, DH1358, and DH1359) that competed with CH235.12 for binding to Env and were derived from the macaque gene ortholog of human VH1-46, IGHV1-105. DH1285 was isolated from NHP7193 and DH1357, DH1358, and DH1359 were isolated from NHP7196 (**Figure 2A and S6A**). DH1357 and DH1358 were clonally related. DH1358 and DH1359 had the same IGHV sequence, but different light chains (**Figure 3A and S8**). DH1285 had a 15 amino acid HCDR3 like CH235.12, whereas the other 3 VH-105 rhesus antibodies had 19 amino acid HCDR3s (**Figure 3A**).

**Figure 3.**
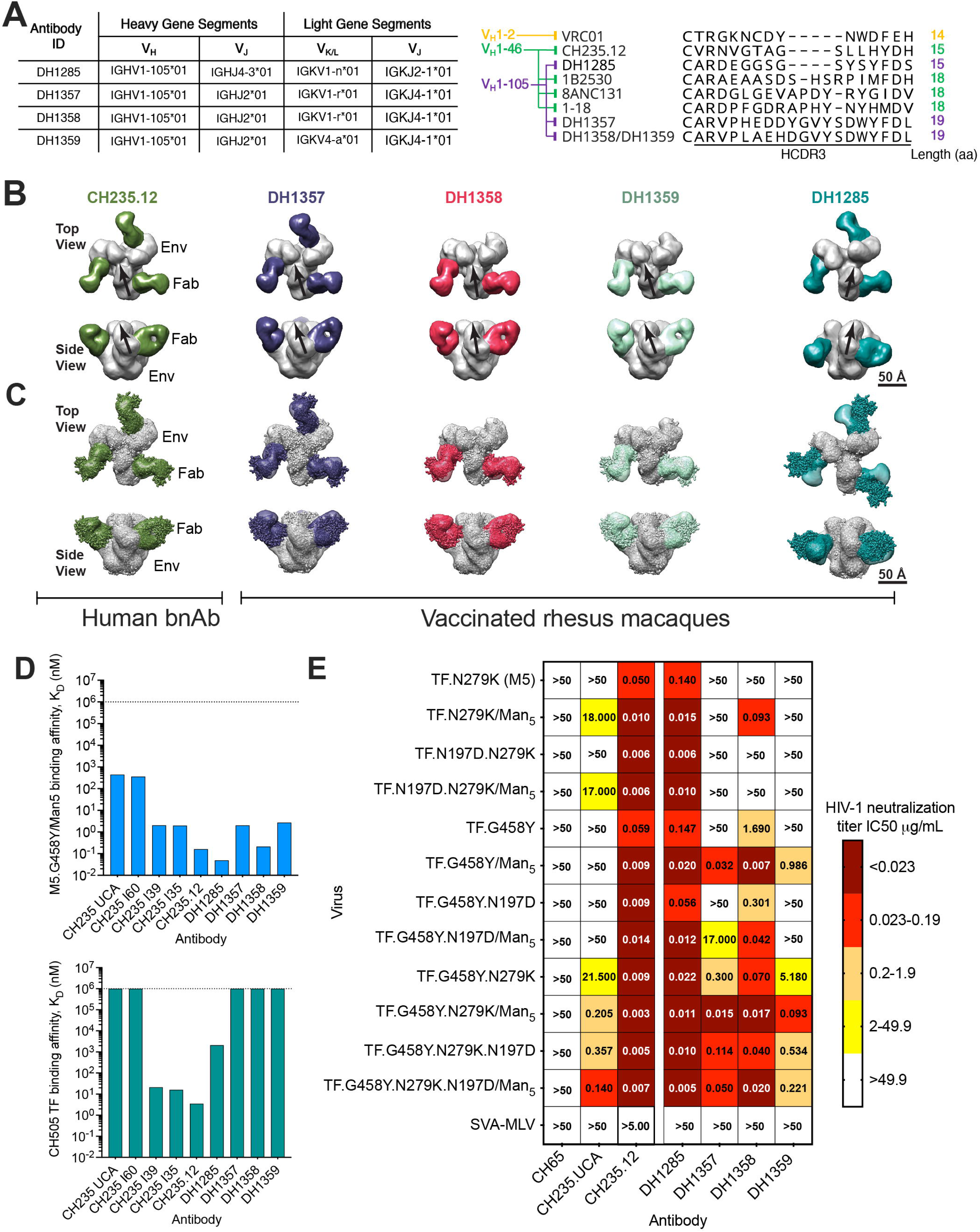
Characterization of Four CH235-like Precursors Isolated from Two Vaccinated Macaques. **(A)** Immunogenetics of putative macaque antibody precursors (left) and HCDR3 alignment compared to human CD4bs bnAbs (right). **(B)** Negative stain electron microscopy shows approach angles for different Fabs binding to the CD4bs of CH505.M5.G458Y. Human CD4-mimicking bnAb CH235.12 is show for comparison. The gp120 axis is indicated by a black arrow. Note the gp120 is rotated in the DH1285 bound Env. (**C**) Superposition of CH235.12 and rhesus Fab in complex with Env. CH235.12 bound to Env is the structure shown in spheres representation. **(D)** Comparison of Fab binding affinity for M5.G458Y (top) or TF (bottom) stabilized gp140 Env trimers. Dashed lines indicate limit of detection. **(E)** Macaque nAb IC50 neutralization titers against CH505 pseudovirus variants with different CH235 enabling substitutions. Macaque nAb titers are compared to the CH235 lineage bnAb putative precursor (CH235 UCA) or bnAb CH235.12.

We performed negative stain electron microscopy (NSEM) to determine whether these antibodies exhibited Env binding modes similar to CH235.12 (**Figure 3B**). The vaccine-induced macaque Fabs DH1357, DH1358 and DH1359 bound to closed trimers and exhibited angles of approach that were highly similar to each other and to the human bnAb CH235.12 (**Figure 3B**). The structure of DH1285 was solved in complex with four different stabilized versions of CH505 TF or M5.G458Y gp140 Env. Three of the structures showed DH1285 bound to partially open Env conformations and one structure with M5.G458Y showed it bound to a closed Env (**Figure 3B and S9**). Although the individual variable loops cannot be resolved by NSEM, the overall density and shape of the partially open Env protomers in the DH1285-bound structure were consistent with an open-occluded Env trimer in which the protomers rotate away from one another to open the trimer, but the V1V2 loops remain in their closed position occluding the V3 co-receptor binding site^34^ (**Figure 3B and S9A-C**). Altogether, the collection of structures showed DH1285 bound to both open and closed Env conformations, demonstrating that DH1285 can bind multiple Env conformations and that a partially open conformation was not required for binding (**Figure S9**). A high-resolution model of the CH235.12 complex fit into the NSEM density maps for DH1357, DH1358 and DH1359 with only small deviations in the constant regions of the Fabs, thus confirming their similarity to CH235.12 (**Figure 3C**). In contrast, when the CH235.12 complex is modeled into the DH1285 structure it can be seen that DH1285 targets the same epitope as CH235.12, but with a different angle of approach that is most apparent when viewed from the top looking down the trimer axis (**Figure 3C**).

Next, we compared the binding affinity of the vaccine-induced antibodies to that of CH235.12 lineage members. DH1285 showed the highest affinity (0.049 nM) for Man_5_GlcNAc_2_−enriched M5.G458Y among all macaque and human antibodies tested (**Figure 3D**). Both DH1358 and CH235.12 bound to Man_5_GlcNAc_2_-enriched M5.G458Y with approximately 0.2 nM affinity (**Figure 3D**). DH1357 and DH1359 Fab affinity for Man_5_GlcNAc_2_-enriched M5.G458Y were most similar to the later intermediates (CH235 I39 and I35) in the CH235.12 lineage (**Figure 3D**). For the CH505 TF Env trimer, CH235 I39, CH235 I35, and CH235.12 had detectable affinity, while DH1285 was the only macaque antibody that bound the CH505 TF Env (**Figure 3D**). The DH1285 affinity (2,070 nM) was 100-fold weaker than CH235 I39 or CH235 I35 for CH505 TF Env (**Figure 3D**). Overall, vaccine-induced rhesus antibodies had extremely high affinities for the vaccine immunogen but were most similar to early intermediate antibodies in the CH235 lineage when binding to a wildtype Env.

We next determined antibody neutralization of a panel of CH505 TF viruses that included combinations of Env modifications that enabled CH235 precursor binding to Env. Of the vaccine-induced antibodies, DH1285 showed the broadest neutralization of the variant CH505 viruses. The neutralization potency of DH1285 closely resembled the potency of CH235.12. DH1358 exhibited the second broadest neutralization, showing a preference for either Man_5_GlcNAc_2_-enrichment or G458Y to be present in the virus. The neutralization profile of rhesus CD4bs Ab DH1359 was most similar to CH235 UCA in that it was still highly dependent on N279K and G458Y. Four of the five viruses DH1359 neutralized had both N279K and G458Y. DH1357 and DH1359 showed similar neutralization patterns, although neutralization was more potent for DH1357. DH1285 and DH1357 were the highest affinity antibodies and were the least dependent on Env modifications that enabled CH235 precursor binding such as N197 glycan removal, N279K, G458Y, and Man_5_GlcNAc_2_ enrichment. Thus, of the four CH235.12-blocking antibodies, the binding phenotypes of DH1285 and DH1357 have progressed the furthest from the precursor stage.

### DH1285 exhibits the canonical CD4 mimicking structure of VH1-46 and VRC01 bnAb classes

To obtain higher resolution definition of DH1285 binding to HIV-1 Env, we determined cryo-EM structures of the DH1285 Fab bound to a stabilized CH505 TF SOSIP Env (**Figure 4A, S10, S11, and Table S2**). The particles in the cryo-EM dataset, picked and sorted using reference-free 2D classification, revealed DH1285 Fabs bound to the Env trimer in the 2D class averages. A heterogeneous mix of particles were observed, from which *ab initio* models were generated and particles were sorted via multiple rounds of heterogeneous refinements, yielding distinct classes showing compositional heterogeneity arising from varied occupancy of the DH1285 Fabs per Env trimer, as well as conformational heterogeneity arising from different relative orientations of the Env protomer. Cryo-EM reconstructions of DH1285 Fab bound to the CH505 Env trimer were resolved to a global resolution ranging from 5-6 Å (**Figure 4A, S10, S11 and Table S2**). Local refinement of the DH1285 Fab interface with the Env yielded a 4.5 Å resolution reconstruction, enabling unambiguous placement of the Fab and resolution of interfacial loops and secondary structures, including the CD4 binding loop, and Loops D and V5 of Env gp120 and the antibody CDR loops (**Figure 4C**).

**Figure 4.**
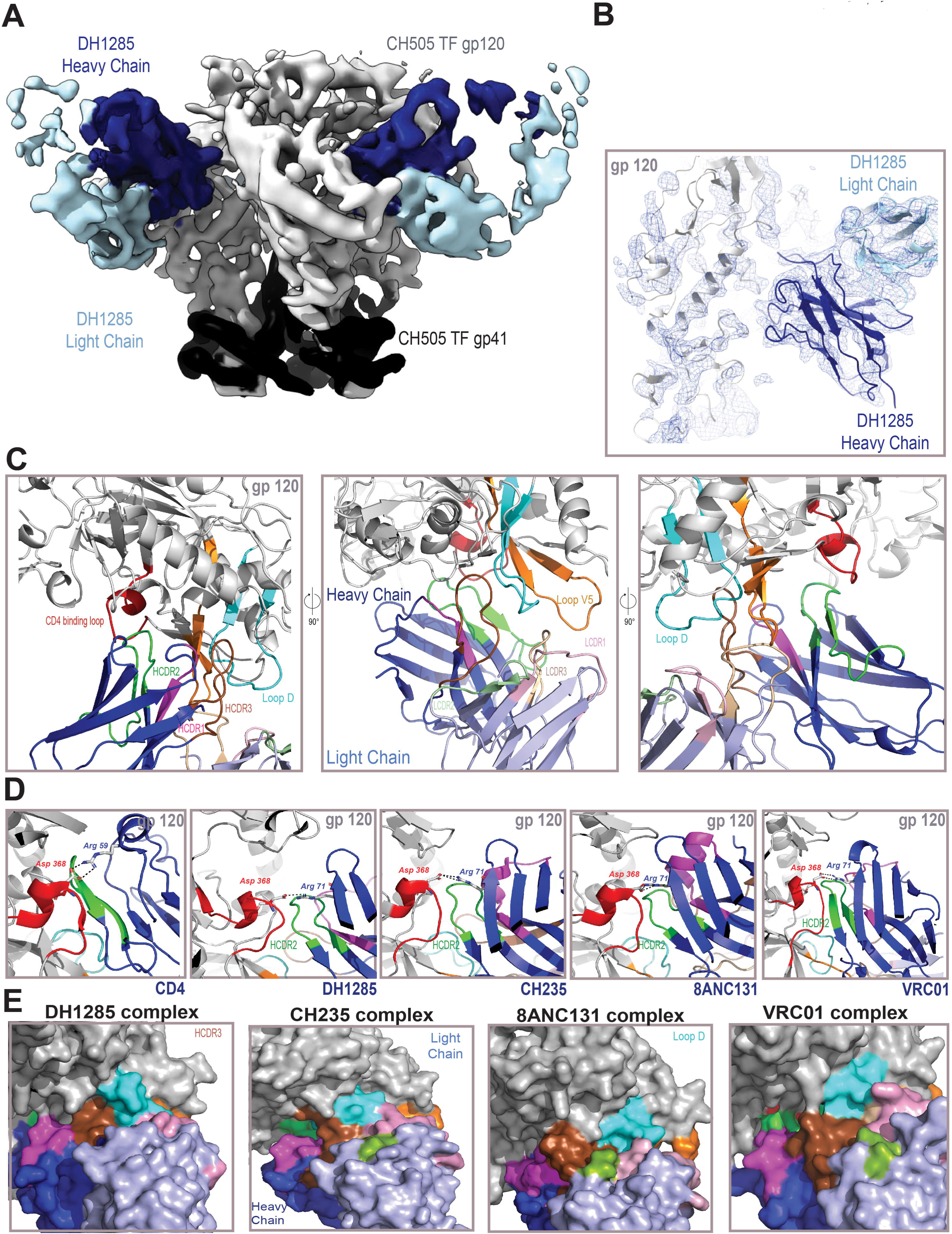
Cryo-EM Structure of DH1285 Bound to CH505 TF Env Demonstrates Antibody Mimicry of CD4 like Human BnAbs. **(A)** Cryo-EM reconstruction of DH1285 Fab (heavy chain in dark blue light chain in light blue) in complex with CH505 TF Env SOSIP (gp120 in light gray, gp41 in black). **(B)** Cryo-EM reconstruction (shown as a blue mesh) from local refinement of the gp120/Fab interface, with underlying fitted model shown in cartoon representation. **(C)** Three views of DH1285 bound to CH505 TF gp120; gp120 colored gray with the CD4 binding loop colored red, Loop D colored cyan, and Loop V5 colored orange. DH1285 heavy chain colored dark blue with HCDR1, HCDR2 and HCDR3 colored magenta, green and brown, respectively. DH1285 heavy chain colored dark blue with HCDR1, HCDR2 and HCDR3 colored magenta, green and brown, respectively. DH1285 light chain colored light blue with LCDR1, LCDR2 and LCDR3 colored pink, light green, and light brown, respectively. **(D)** Interactions of the CD4 binding loop (red) shown with, (from left to right) CD4, DH1285, CH235, 8ANC131, and VRC01. Residue Asp 368 in gp120, Arg 71 in the antibody heavy chains and Arg 59 in CD4 are shown in stick representation. The salt bridge between Asp 368 and Arg 71 in the antibodies or between Asp 368 and Arg 59 in CD4 are shown as dashed lines. **(E)** Surface representation showing the interactions of gp120 Loop D (cyan) with the bound antibody. Antibody heavy chain is shown in dark blue and light chain in light blue; HCDR1, HCDR2 and HCDR3 colored magenta, green and brown, respectively

Human CD4bs bnAbs that mimic CD4 bind with stereotypical paratope structures that resemble portions of CD4^9,13,15^. Specifically, each human bnAb has a HCDR2 in a beta strand conformation that contacts the CD4 binding loop, and an arginine at position 71 in the V_H_ that forms a salt bridge with aspartic acid at position 368 in Env (**Figure 4D**). DH1285 antibody interacts with gp120 using both heavy and light chain complementary determining regions (CDRs). The DH1285 HCDR2 interacts with the CD4 binding loop of gp120 mimicking the gp120-CD4 receptor interaction in this region, with residue Asp168 of the CD4 binding loop positioned to make a salt bridge with Arg 71 in the antibody framework region 3. Therefore, DH1285 exhibits the conserved structural and interactive signatures of the VH1-2 and VH1-46 CD4-mimetic antibodies.

The DH1285 HCDR3 and LCDR3 regions contacted both gp120 Loop D and Loop V5, respectively (**Figure 4C and 4E**), with the DH1285 HCDR3 interaction with Loop D most closely resembling that of CH235.12. The HCDR3 also contacted structural elements in the gp120 inner domain with the gp120 Trp96 side chain stacking against the HCDR3 tip. Loop D also contacted the light chain LCDR1 and LCDR3 regions as well as Trp 50 of the HCDR2, making Loop D a key interactive region that contacted spatially separated regions of the antibody. While LCDR2 showed no direct contact with the epitope, it contacted the LCDR1 and LCDR3 loops and may play a role in stabilizing the conformations of these paratope loops and influencing their presentation. Comparison of gp120-CD4 (PDB: 1GC1), gp120-VRC01(PDB: 3NGB), gp120-8ANC131(PDB: 4RWY), gp120-CH235(PDB: 5F9W) on gp120-DH1285 complex reveals their relative orientations and highlights the similarity of their interactions centered on HCDR2-CD4 binding loop (**Figure 4E**). Taken together, the structural data confirm that antibody DH1285 binding to HIV-1 Env recapitulates the key structural signatures that are hallmarks of VH1-2 and VH1-46 germline derived CD4-mimetic HIV-1 bnAbs.

### DH1285 and CD4-mimicking human bnAbs share molecular immunology features for Env binding

The combination of high-resolution structures of VRC01 and CH235 precursors have identified W50, R71, and N58 as germline V_H_ sequence-encoded amino acids that mediate contact within the CD4bs on Env^9,15, 35–37^. These amino acids are postulated to be the genetic basis for why specific IGHV gene segments are used by CD4 mimicking bnAbs. Also, V_H_ amino acids at position 54 substantially affect binding affinity of CD4 mimicking antibodies by inserting their side chains into the cavity on Env usually occupied by F43 in CD4, termed the Phe43 cavity^38^. We compared the V_H_ sequence and structure of DH1285 to known VRC01 class and VH1-46 class bnAbs, identified that the same amino acids at positions 50, 54, and 71 as VH1-46 or VRC01 class antibodies. At position 58, DH1285 had a K58H substitution that differed from the human bnAbs examined (**Figure 5A and B**). In addition to the R71 described above, the DH1285 HCDR2 was positioned similarly to human CD4bs bnAbs such that the DH1285 VH W50 and Y54 contacted the CD4bs in a manner akin to VRC01, NIH45-46, CH235 UCA, 1-18, and 3BNC117 (**Figure 5A and B**). Alanine substitution of W50, Y54, and R71 showed that not only were they shared amino acids, but they also were required for optimal binding to CH505 TF Env trimer and Man_5_GlcNAc_2_-enriched CH505 TF neutralization (**Figure 5A and C**). Alanine scanning mutagenesis of the entire HCDR2 further showed that N52, P52a, and N56 were required for optimal Env binding and neutralization of the same protein and virus, respectively (**Figure 5C**). Given the proximity of R73 to R71, we also substituted alanine at this position, although it should be noted that R73 is a result of somatic mutation. R73A substitution moderately reduced CH505 TF neutralization (**Figure 5C**). Binding and neutralization of Man_5_GlcNAc_2_-enriched M5.G458Y was only affected by R71 substitution, suggesting the Env engineering to improve affinity for CH235-like antibodies compensated for most single amino acid substitutions (**Figure S12A and S12B**).

**Figure 5.**
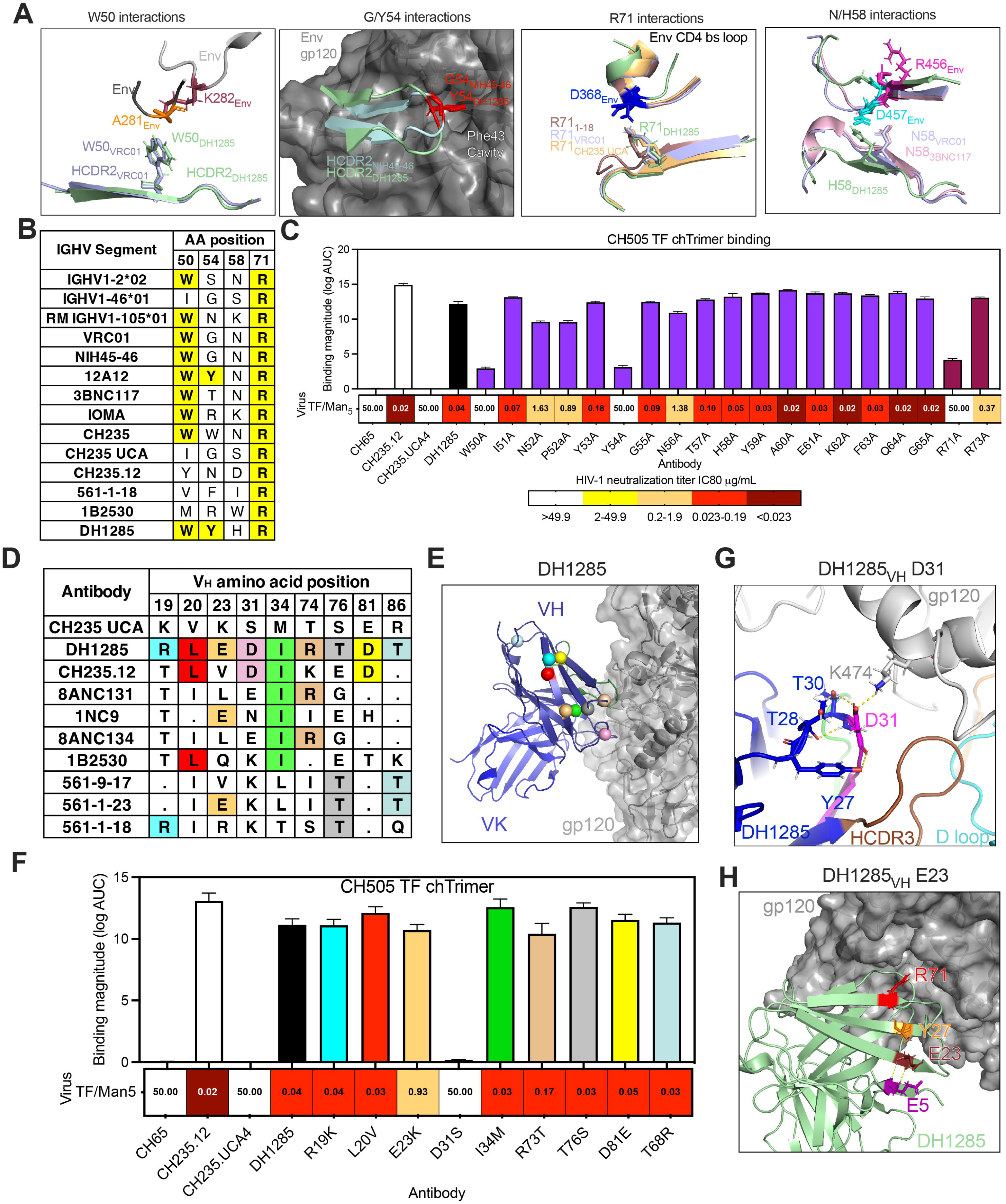
Molecular Features Are Conserved Among DH1285 And Human CD4-Mimicking BnAbs. **(A,B)** DH1285 has the key amino acids for interaction with the CD4bs (antibody amino acids 50, 54, 58, and 71) that were previously identified in human CD4-mimicking bnAbs. **(A)** Antibody amino acid interactions by human bnAbs and DH1285 with Env are superimposed (PDBs:6UDA, 5V8M, 6UDJ, 5WDU, 6V8X). Of note are the interactions between the D loop and W50, the Phe43 cavity and Y54, the CD4bs loop and R71, and the V5 loop and H58. **(B)** Comparison of residue identity at key sites listed in (**A**) and their conservation among V_H_ gene segment-restricted CD4bs bnAbs. Relevant human and rhesus germlines are shown in the first three rows. Shared amino acids with DH1285 are shown in bold and highlighted yellow. **(C)** Binding and neutralization activity of the DH1285 HCDR2, R71, and R73 alanine mutants. CH505.TF Env trimer binding mean log AUC of three independent experiments and standard error of mean are shown. Man_5_GlcNAc_2_-enriched CH505 TF pseudovirus neutralization IC_80_ titers for the DH1285 alanine mutant antibodies are shown under the binding magnitude graph. **(D)** V_H_ amino acids encoded for by somatic mutations that are conserved between DH1285 and VH1-46 bnAbs are shown in colored boxes. Boxes with periods indicate amino acids that are identical to the CH235 UCA. **(E)** Structural location of conserved DH1285 and VH1-46 bnAb mutations relative to the HIV-1 gp120 interface. The conserved amino acids are shown as spheres within the DH1285 variable region and are color-coded as in **(D)**. **(F)** CH505 TF Env trimer binding by antibodies with conserved amino acids reverted to the VH1-46 germline amino acid. Binding magnitude is shown as the mean logAUC from triplicate experiments with standard error of the mean. The heatmap under the bar graph shows the neutralization IC_80_ titer for the same mutant antibodies against Man_5_GlcNAc_2_-enriched CH505 TF pseudovirus. Neutralization titer is color-coded as in **C. (G)** DH1285 cryo-EM structure showing D31_DH1285_ contacts with K474_Env_. **(H)** DH1285 cryo-EM structure showing positioning of E23_DH1285_ relative to other framework region residues with potential structural coordination of the positioning of the R71 for contact with the CD4bs loop.

DH1285 also shared identity with nine amino acids that resulted from somatic mutation in VH1-46 class bnAbs (**Figure 5D and S12C**). While some of these amino acids may be shared due to hotspots for somatic mutations, we hypothesized that a subset of the nine mutations were shared because they improved Env binding for this class of antibody. Examination of the CH235 UCA and DH1285 Fabs in complex in Env supported this hypothesis since four of the amino acids were located at the antigen-antibody interface (**Figure 5E**). We changed the shared amino acids to the germline VH1-46 amino acids to determine whether the shared amino acid contributed to antibody binding or neutralization (**Figure 5F, S12D, and S12E**). Six single amino acid changes had no effect on binding or neutralization (**Figure 5F, S12D, and S12E**). R73T and E23K showed a 4-fold and 23-fold decrease in Man_5_GlcNAc_2_-enriched CH505 TF neutralization potency respectively, but both changes caused minor reductions in recombinant Env binding (**Figure 5F, S12D, and S12E**). In contrast, D31S substitution completely abrogated CH505 TF gp140 trimer binding and Man_5_GlcNAc_2_-enriched CH505 TF neutralization. The structural basis for the dependence on D31 was that D31 interacted with K474 in the CD4bs of HIV Env (**Figure 5G**). It also coordinated the overall HCDR1 conformation through interactions with Y27, T28, and T30 in the first framework of DH1285 (**Figure 5G**). DH1285 E23 did not mediate direct contact with Env but coordinated the beta sheet conformation that ultimately positions R71 to interact with the CD4bs loop (**Figure 5H**). Altogether, DH1285 has functional canonical germline-encoded and somatic mutation-encoded amino acids found in CD4 mimicking human bnAbs.

### Env engineering for CH235 UCA binding enables binding by the DH1285 UCA

The M5.G458Y immunogen was designed to bind to the precursor of CH235 lineage. However, it is unknown how well it can bind to orthologous precursors in nonhuman primates that are not exactly the same sequence as the CH235 UCA. To infer the UCA antibody that gave rise to the DH1285 lineage, we performed two independent MiSeq next-generation sequencing (NGS) runs of macaque VH1 and VK1 regions of peripheral blood B cells. We used Cloanalyst and Partis to assign clonality to the recovered heavy chain and light chain sequences and inferred a DH1285 UCA. We ran IgDiscover using our MiSeq sequences and found no evidence for a new IGHV1-105 allele being present in macaque 7193^39^. Thus, the UCA is composed of previously known V gene segments. We examined the binding affinity of the UCA for the immunogen designed to target CH235-like unmutated antibodies. The apparent affinity of the DH1285 IgG was 36.8 nM, which was 8-fold weaker than the CH235 UCA apparent affinity of 4.5 nM (**Figure 6A, S13A**, **and S13B**). A fast on-rate between the antigen and B cell receptor is associated with the ability of the B cell receptor to signal^40^. We found the on-rate between the CH235 and DH1285 UCA were similar, differing by only 5-fold (6.86E4 and 1.35E4 respectively) (**Figure 6A and S13B**). Neither UCA bound to the CH505 TF (**Figure 6A**). Thus, antibody precursors with 8-fold weaker apparent binding affinity and 5-fold slower on-rates than the CH235 precursor can be engaged and expanded by the engineered Env M5.G458Y.

**Figure 6.**
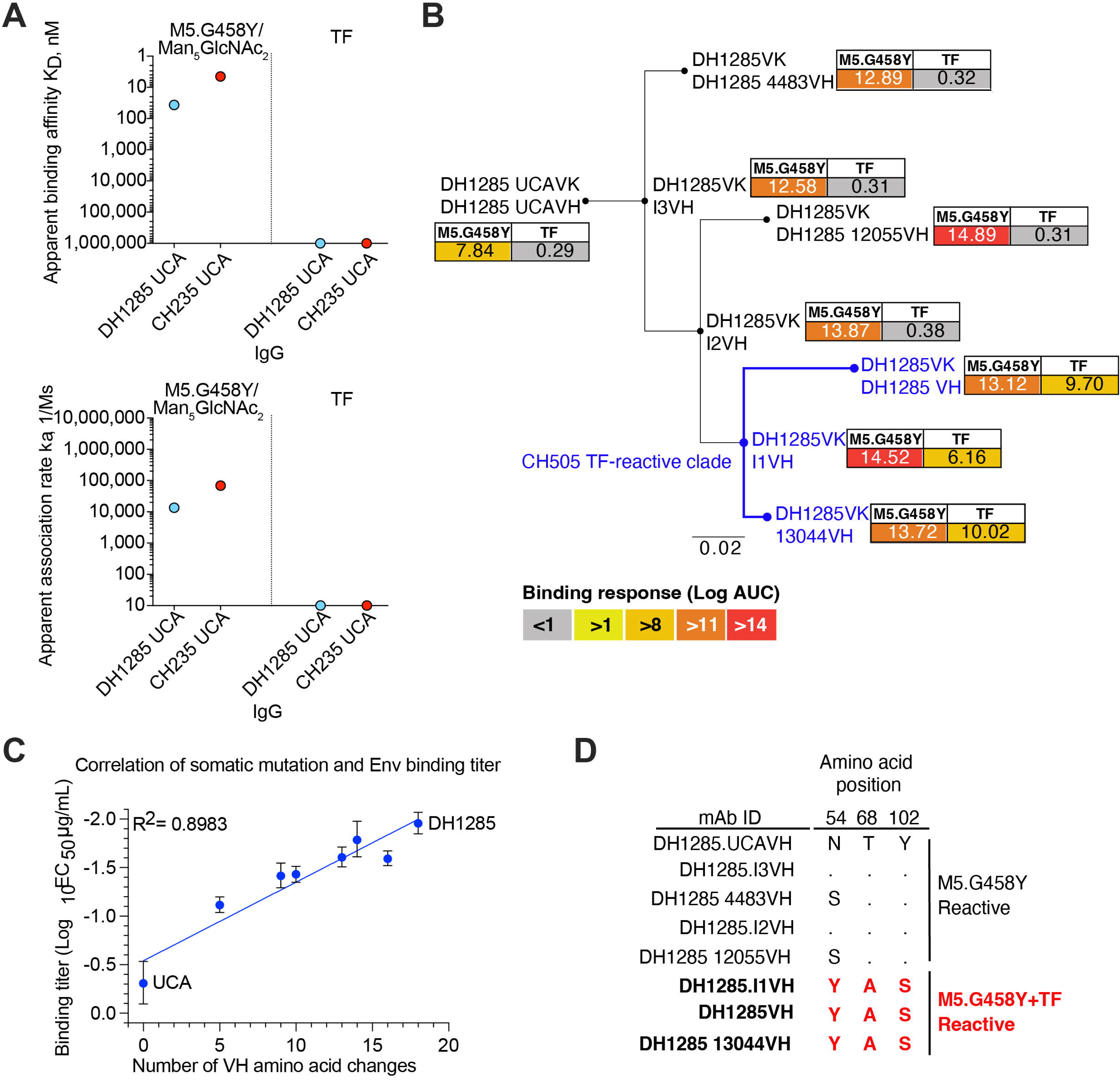
DH1285 Ontogeny Shows the Putative DH1285 Precursor Antibody Binds the Vaccine Immunogen and the Emergence of CH505 TF Env Recognition. **(A)** (Top) Apparent binding affinity and (bottom) association-rate of DH1285 UCA and CH235 UCA IgG binding to Man_5_GlcNAc_2_-enriched CH505.M5.G458Y or TF Env trimer. **(B)** Phylogenetic tree of DH1285 clonally-related VH sequences. Terminal nodes show observed sequences with internal nodes indicating inferred intermediates. Binding magnitude for Man_5_GlcNAc_2_-enriched CH505.M5.G458Y and CH505.TF gp140 was determined for each DH1285 clonal member when paired with the DH1285 light chain. Antibodies that react with CH505 TF Env trimer are colored blue. Values are the mean of 2 independent experiments. **(C)** Spearman’s correlation of M5.G458Y (blue) Env trimer log_10_ EC_50_ binding titer and the number of DH1285 VH amino acid substitutions. **(D)** Amino acid residues at three sites that mutate when CH505 TF Env trimer binding is first observed in the DH1285 lineage.

To understand how the heavy chain evolved from the UCA, we inferred heavy chains within the DH1285 lineage using NGS sequences that were observed in two independent sequencing runs (**Figure 6B and S13A**). The NGS-derived DH1285 lineage V_H_ sequences were paired with the DH1285 light chain and produced as recombinant IgGs for phenotypic characterization. The DH1285 UCA bound to Man_5_GlCNAc_2_-enriched M5.G458Y Env trimer by ELISA. The binding magnitude for Man_5_GlCNAc_2_-enriched M5.G458Y Env trimer increased by less than 2-fold in logAUC as the DH1285 clone continued to evolve, with the major increase in binding magnitude occurring at the first intermediate antibody (I3; **Figure 6B and S13C**). Overall, binding magnitude to Man_5_GlCNAc_2-_enriched M5.G458Y Env trimer increased as the V_H_ region acquired more amino acid changes (**Figure 6B**). Showing the importance of the Env modifications that promote CH235 precursor binding, the DH1285 UCA showed negligible binding to CH505 TF Env trimer lacking these modifications (**Figure 6B**). Thus, the modification of Env to enable CH235 precursor binding also enabled vaccine-induced rhesus CD4bs DH1285 UCA binding. Within the DH1285 antibody clone, CH505 TF Env reactivity was only detected by DH1285, DH1285 I1 and DH1285.13044, which were all from the same clade of the phylogeny (**Figure 6B**). In contrast to M5.G458Y binding, CH505 TF binding did not strongly correlate with number of antibody amino acid changes suggesting the requirement for specific critical amino acid substitutions rather than a particular number of amino acid changes (**Figure S13C, and S13D**). Comparison of the VH sequences of the CH505 TF-reactive antibodies to the other clonally related V_H_ sequences that lacked CH505 TF binding showed amino acid changes N54Y, T68A, and Y102S (**Figure 6D and S13E**). Alanine substitution of Y54 demonstrated that this tyrosine was required for CH505 TF Env binding (**Figure 5C**). Hence, DH1285 UCA BCR precursor engaged the engineered immunogen in vivo after macaque immunization and affinity matured to include known key somatic mutations such as N54Y, that enabled interaction with wildtype envelope.

## DISCUSSION

Here we demonstrate here that an HIV-1 immunogen designed to engage the VH1-46 class CH235 bnAb lineage selects for similar BCR in outbred nonhuman primates defined by structural mode of Env binding, amino acid mutations acquired, and usage of the rhesus macaque ortholog of human VH1-46 gene. Thus, BCRs that can give rise to a VH1-46 bnAb as antibodies derived for an orthologous immunoglobulin heavy chain variable gene segment, contains the critical R71_VH_, CD4bs-dependent binding, and HCDR2-mediated binding to the HIV-1 Env CD4 binding loop--criteria selected based on immunogenetic and structural studies of the VH1-46-derived CD4bs bnAbs^6,11,13,14^. These characteristics are also hallmarks of most if not all CD4-mimicking bnAbs including VRC01 and 3BNC117. Remarkably, our structural and immunogenetic studies demonstrate that DH1285 possessed all four of these criteria for classifying it as a VH1-46-like bnAb precursor. In addition to R71_VH_, the VH1-105-derived antibodies also possessed W50_VH_ in HCDR2, which is another shared trait of CD4-mimicking CD4bs antibodies such as VRC01 and 3BNC117^35–37^. Interestingly, CH235 is thought to somatically mutate to encode W50, although the germline gene of the HIV-infected individual was not sequenced to confirm W50 was not present in its allele of IGHV1-46^13^. The shared amino acids encoded by human IGHV1-46 and macaque IGHV1-105 highlight the genetic similarities between humans and macaques. It has been posited that macaques lack the necessary germline encoded gene segments to express CD4-mimicking neutralizing antibodies. To the contrary, we find here that indeed macaques can make CD4bs antibodies with sufficient germline amino acids that form the initial, critical contacts made with Env. Although we only administered an Env designed to engage and expand germline antibodies similar to the VH1-46 bnAbs from humans, we found that solated BCRs that were expressed as recombinant antibodies had begun the process of affinity maturation to include residues similar to the VH1-46 bnAbs. Thus, the engineered HIV-1 Env immunogen was successful in eliciting the desired target CD4bs antibodies and began to select for an affinity maturation pathway similar to human VH1-46 class bnAbs. It should also be noted that the neutralizing CD4bs antibodies elicited in the macaques could be detected in the serum. Therefore, a substantial portion of the serum neutralizing antibody response was directed to the CD4bs.

Three vaccine-induced IGHV1-105-derived antibodies showed highly similar angles of approach to CH235. A fourth IGHV1-105-derived antibody showed a distinct angle of approach but same contact site on Env. This result is significant since structural studies of the CH235 lineage has demonstrated that the CH235 angle of approach is determined at the UCA stage of the lineage and is unchanged during affinity maturation^13^. This observation suggests that human VH1-46 bnAb precursors will also need to adopt the correct binding angle and orientation at their precursor stage. For macaque or human antibodies whose angles of approach differ from CH235.12, it is uncertain whether they can evolve to bind to Env in a manner similar to CH235. The observation that the macaque antibodies described here have the appropriate initial binding mode is encouraging for trying to select for higher affinity-matured bnAb intermediates with greater neutralization breadth.

The germline targeting approach where immunogens are designed to interact with high affinity to one or a few known bnAb antibody precursors is a major strategy for HIV-1 vaccine development ^5,23–25^. The G001 trial has provided proof-of-concept for targeting putative VRC01-class CD4bs antibodies in humans although structural confirmation of HCDR2 binding of precursors was not reported^41^. VRC01 affinity maturation includes selection of rare deletions in the light chain that may prove difficult for vaccine elicitation ^42^. Thus, vaccine designs targeting additional CD4bs bnAb classes are warranted. The CH235 lineage is an advantageous vaccine target since it lacks rare nucleotide deletions or an unusually short light chain third complementarity determining region observed in the VRC01 lineage^10,13,28^. We show here that such CD4bs bnAb precursor B cells can be expanded with the Man_5_GlcNAc_2_-enriched M5.G458Y immunogen. Although the immunogen was designed based on the human CH235 precursor antibody, it was able to bind with nanomolar apparent affinity to a rhesus macaque inferred germline CD4bs IgG antibody. This cross-species germline targeting of CD4bs bnAbs shows the immunogen is quite adept at interacting with these precursors, as well as shows the validity of using rhesus macaques as a model for development of vaccine immunogens that can target VH1-46 CD4bs bnAb lineages. This precise targeting of CD4bs antibodies may explain why we observe extremely high titers of CD4bs-dependent neutralizing antibodies exhibiting the CH235 neutralization signature in the serum. Overall, this study supports the notion that immunogens designed based on a known antibody lineage can elicit similar antibody lineages in unrelated individuals or even different primate species. The HVTN309 Phase I trial will test this hypothesis in humans by administering M5.G458Y envelope nanoparticles with ionizable LNP as an adjuvant to elicit VH1-46 class bnAb precursors.

Due to the protracted affinity maturation process for bnAb development, a single Env is not expected to select for affinity maturation from germline antibody to bnAb ^23,43^. However, the antibodies isolated after the priming immunization can inform the next steps in sequential vaccine design. We found here that the processed glycan at position 197 on CH505 TF Env impeded DH1285 neutralization of natively glycosylated CH505 TF pseudovirus. Vaccine immunogens that select for affinity matured DH1285 that can accommodate processed glycan at position 197 would be a logical next step in sequential vaccine design. There are clues from the VH1-46 bnAb class as to how affinity maturation can accommodate this glycan. The CH235 lineage makes improbable K19T, E81D and T70Y amino acid substitution in its heavy chain precisely where the glycan juxtaposes the antibody framework region. Vaccine immunogens that select for such BCR mutations in DH1285 antibodies may enable it to neutralize natively glycosylated wildtype viruses. In the initial antibody-virus coevolution study of the individual who generated the CH235 lineage, Envs with elongated and glycosylated fifth variable regions arose over time^27^. Selecting for antibodies that can accommodate changes in the fifth variable region of the CD4bs is hypothesized to be one pathway towards developing neutralization breadth.

Adjuvants play a critical role in the vaccine-induced immune response. In a previous study in mice, we showed that LNP were potent adjuvants when mixed with protein immunogens and induced robust Tfh responses^29^. Here, LNP were potent inducers of serum antibody responses with as little as two immunizations and induced robust Tfh and GC B cell responses after three immunizations. While Env trimers have been viewed as less immunogenic than nanoparticle vaccines^44–46^, lipid nanoparticles mixed with HIV-1 Env trimers were able to elicit potent neutralizing antibodies.

We acknowledge the study limitations are that it is difficult to determine whether a particular antibody will develop into a bnAb based on its early stages of development. For previous germline-targeting HIV vaccines, the bnAb precursors are usually non-neutralizing^41^, however the bnAb precursors elicited here demonstrate nanogram per milliliter IC80 neutralization titers. Thus, the neutralization potency combined with the favorable genetic and structural traits make development into a CD4bs bnAb a plausible outcome. We also acknowledge one limitation is that we immunized six times which is not conducive to heterologous prime boost sequential vaccines. Lastly, we tested only one dose of LNP as an adjuvant. The adjuvanting effect of other doses of LNP still needs to investigated in future studies.

In summary, proof of concept for VH1-46-type, CD4 mimicking, CD4bs bnAb precursors has been achieved in outbred non-human primates, providing evidence for the possibility that fully affinity-matured bnAbs will be able to be induced in both rhesus macaques and in healthy humans by judicious choice of a series of boosting immunogens that can sequentially select for functional improbable mutations necessary for full bnAb potency and breadth.

## MATERIALS AND METHODS

### Animals and Immunization

Indian-origin rhesus macaques were housed in AAALAC-accredited facilities and all veterinary and study procedures performed in accordance with Duke University IACUC-approved protocols and Bioqual (Rockville, MD) standard operating procedures. All macaques were 2 years and 4 months old at the start of the study, and weighed between 5.2 and 6.8 kg at the end of the study. The study began with 4 macaques, but one female macaque, NHP 7194 was removed from the study for animal medicine reasons, leaving 3 (2 female and 1 male) macaques that completed the study. The first 5 immunizations were spaced every four saweeks with a final immunization 14 weeks after the 5^th^ immunization. Vaccine was administered intramuscularly by injecting 750 µL of protein plus adjuvant mixture in both the left and right quadriceps (total 1.5 mL injection per animal per immunization). Each immunization consisted of 100 µg CH505.M5.G458Y Man_5_GlcNA_c2_-enriched Env trimer formulated with nucleoside-modified mRNA encoding luciferase encapsulated in lipid nanoparticles from Acuitas. Throughout the duration of the study, whole blood and serum were drawn on the day of vaccination and one and/or two weeks post-vaccination.

### HIV Env Protein Production

Previously described chimeric CH505 TF and CH505.M5.G458Y SOSIP gp140 Env proteins^15,21^ were stabilized with E64K and A316W stabilizing mutations^47^. The codon-optimized genes expressing the trimers were encoded by the VRC8400 vector.

For expression, Freestyle 293F (ThermoFisher) or 293S GnT1^-^ cells were diluted at the time of transfection to 1.25 x 10^6^ cells/mL with fresh Freestyle293 (ThermoFisher) media in 500 mL batches. Co-Transfection was performed with plasmid DNA (650 µg SOSIP trimer plasmid and 150 µg furin expressing plasmid per 1L of culture volume) complexed with 293fectin in OPTI-MEM. After 6 days, cell cultures were harvested by centrifugation of the cells for 60 minutes at 4000 rpm on a Sorvall table top centrifuge. Supernatant was filtered through 0.8 µm filter and concentrated to approximately 100 mL with Vivaflow 200 cassettes (Sartorius) with a 30 kDa MWCO. Concentrates were again filtered to 0.8 µm, then purified with positive selection on a 10 mL affinity column containing PGT145 conjugated to CnBr-activated Sepharose (Cytvia), buffered with PBS. Following loading and washing, Env trimers were eluted using 3M MgCl_2_, and immediately equilibrated with 5 volumes of 10 mM tris, pH8. The eluate was filtered to 0.2 µm and concentrated to 2 mL with a centricon-70 10 kDa MCWO. If biotinylated (for immunochemistry), Avi-tagged Env proteins at 25 µM were dialyzed into Tris pH8 and incubated for 5 hours with mild agitation at 30 °C using the BirA biotin-protein ligase reaction kit (Avidity LLC), then reconcentrated prior to size-exclusion chromatography. Two milliliters of concentrated protein was purified on a Superose6 16/600 column (Cytvia) in 500 mM NaCl buffer with 10 mM tris, pH 8 to isolate trimeric protein. All chromatography steps were conducted on a an AKTA Pure (Cytvia). The trimeric pool of protein fractions was pooled, filtered to 0.2 µm, and snap frozen for long-term storage at −80 °C.

Recombinant gp120s were produced via transient transfection using Freestyle 293F (ThermoFisher) or 293S GnT1^-^ cells and 293Fectin using the same conditions as SOSIP gp140s. Following five days of culture, cells were harvested via centrifugation and filtered to 0.8 µm. Cell-free supernatant was concentrated using a 10 kDa MWCO Vivaflow 50 (Sartorius). The concentrate was mixed with *Galanthus nivalis* lectin agarose resin (Vistar Labs) incubated overnight at 4 °C. The resin beads were repetitively pelleted via centrifugation and washed twice by resuspension with MES buffer, before protein was eluted with methyl-_α_-pyranoside. Monomeric protein was purified using Superdex200 (GE Healthcare) size-exclusion chromatography column on an AKTA Pure (Cytvia).

### Site-Directed Mutagenesis

Antibody heavy chain plasmids were mutated using the QuikChange Lightning (Agilent) kit, following manufacturer recommended reaction conditions. Oligos introducing site-specific mutations were designed with the QuikChange Primer Design Tool (Agilent), synthesized and purified via standard desalting by Integrated DNA Technologies. Cloning was performed in XL-10 gold cells and sequence integrity was determined with Sanger sequencing (Azenta) and subsequent DNA alignment using Geneious (BioMatters).

### Recombinant Antibody Production

Monoclonal antibodies and Fabs were encoded in heavy and light chain plasmids and co-transfected into Expi293F (293i) cells using expifectamine. Briefly, 293i cells were diluted to 2.5 x 10^6^ cells/mL with fresh Expi293 media in 100 mL batches and incubated for 4 hours prior to transfection. Fifty micrograms of each plasmid was mixed with expifectamine in OPTI-MEM I, then added to the transfection culture. Expifectamine kit enhancers were added approximately 16-18 hours post-transfection and cultures were incubated for 5 days before harvest. The cultures were centrifuged at 4000 rpm in a Sorval table top centrifuge and supernatant was filtered through 0.8 µm filter. The cell-free supernatant was incubated overnight with either protein A resin (ThermoFisher) for IgG1 or LambdaFabSelect or KappSelect resin (Cytvia) for lambda or kappa containing Fabs, respectively at 4°C. The bead slurry was centrifuged at 1200 rpm for 10 minutes, then aspirated before washing via gravity filtration with 20 mM tris (pH 7) and 350 mM NaCl buffer and subsequently eluting with 2.5% acetic acid. The eluate was neutralized with Trizma (pH 8), then buffer exchanged through repetitive centrifugation in a Vivaspin Turbo-15 concentrator with 25 mM citrate and 125 mM NaCl buffer (pH 6) before final storage at −80 °C.

### AIM assay measuring TFH responses

Env-specific TFH cells were quantified using an Activation-Induced Marker Assay^32,33^, using an overlapping peptide pool spanning CH505 TF gp140 to stimulate lymph node cells *ex vivo*. TFH cells were identified by flow as viable lymphocytes that were CD4^+^ CD8^-^ CXCR5^hi^ PD1^hi^. Env specificity was measured as the frequency of TFH cells co-expressing OX40 and CD25 after *ex vivo* stimulation with an Env peptide pool, after background subtraction of frequency in unstimulated conditions.

### Enzymatically-Digested Antibody Fabs

Antibody Fabs were generated by papain digestion using the Pierce™ Fab Preparation Kit (ThermoFisher Catalog No: 44985). The manufacture’s protocol was followed except IgG antibodies were dialyzed into PBS for 2h prior to digestion and IgG was digested for 16-18 h. SDS-PAGE and Coomassie staining was used to examine the digestion of IgG into Fabs. Fabs were run through a Superdex200 10/300 column in 25mM Citric Acid with 125mM NaCl (Cytiva) to remove any aggregates. Fabs were stored frozen at −80 °C in 25mM Citric Acid 125mM NaCl pH 6 Buffer.

### HIV-1 neutralization assays

Neutralizing antibody assays were performed with HIV-1 Env-pseudotyped viruses and TZM-bl cells (NIH AIDS Research and Reference Reagent Program contributed by John Kappes and Xiaoyun Wu) as described previously^48,49^. Neutralization titers are the reciprocal sample dilution (for serum) or antibody concentration in µg/mL (for mAbs) at which relative luminescence units (RLU) were reduced by 80% or 50% (ID80/IC80 and ID50/IC50 respectively) compared to RLU in virus control wells after subtraction of background RLU in cell only control wells. Serum samples were heat-inactivated at 56 °C for 30 minutes prior to assay.

### Fab-Env complex formation, negative staining, and data analysis

For polyclonal serum Fabs, ∼1 mg of polyclonal Fabs at ∼10 mg/ml were mixed with 20 µg of M5.G458Y Env trimer and incubated overnight at 4 °C. To remove excess unbound Fab, the mixture was separated by size exclusion chromatography on a Superose 6 Increase 10/300 column and fractions eluting at the expected volume for the complex were combined and concentrated with a 100-kDa molecular weight cutoff spin concentrator to a nominal trimer concentration of ∼1 mg/ml. Concentrated sample was then diluted to 0.4 mg/ml with HEPES-buffered saline (HBS), containing 150 mM NaCl, 20 mM HEPES, pH 7.4, augmented with 8 mM glutaraldehyde for crosslinking and incubated for 5 minutes at room temperature. Unreacted glutaraldehyde was quenched by addition of 1 M Tris, pH 7.4 to a final Tris concentration of 80 mM. As needed, quenched samples were diluted to 0.1-0.2 mg/ml with HBS augmented with 5% glycerol or applied without dilution to glow-discharged carbon films on 300 mesh copper EM grids for negative staining. After blotting excess sample, samples were stained for 1 minute with 2% uranyl formate, blotted and allowed to air dry.

For monoclonal Fabs, 36 µg of Fab was mixed with 10 µg M5.G458Y Env trimer in a total volume of 100 µl of HBS and incubated overnight at 4 °C. Fab-trimer complexes were then diluted with 400 µl of HBS augmented with 10 mM glutaraldehyde, incubated for 5 minutes at room temperature, and quenched by addition of 1 M Tris to 80 mM final concentration. Quenched samples were then concentrated with a 100-kDA molecular weight cutoff spin concentrator, which allows excess unbound Fabs to pass the filter and retains the Fab-trimer complex. Concentrated sample was then diluted and negatively stained as described above.

Negatively stained grids were examined on a Philips EM420 electron microscope operating at 120 kV, 49,000x nominal magnification, and ∼0.5 µm defocus. Images were acquired on a 76-megapixel CCD camera, corresponding to a nominal calibration of 2.4 Å/pixel. Datasets were typically ∼100 or ∼500 images for monoclonal or polyclonal samples, respectively. Image analysis was performed with standard protocols in Relion 3.0 ^50^, beginning with automated particle picking, followed with two rounds of 2D classification/selection, and then by 1-2 rounds of 3D classification/selection to discard junk particles and select Fab-bound trimer particles. For monoclonal samples, particle stacks from well-resolved 3D classes were chosen and final 3D refinements with post-processing performed. For polyclonal samples, the initial 3D classifications were used to estimate the epitope occupancy for each polyclonal sample (Figure S4), and the particles from all Fab-bound classes were combined and further analyzed as described below.

Subsequent analysis of polyclonal samples follows that of Antanasijevic et al.^51^ with slight modifications. Briefly, the entire particle stack was refined to a single structure with C3 symmetry imposed and then symmetry expanded. The symmetry expanded particle stack was subjected to focused 3D classification without alignment using a 100-Å diameter spherical mask centered on the Fabs bound to the CD4bs of one protomer. Classes showing a Fab-like density were selected and their particles combined and subjected to an unmasked C1 refinement with local angular searches only. The location of the 100-Å mask was adjusted as needed to contain all Fabs observed, and a second round of masked 3D classification without alignment performed. Fab-containing classes were selected and their combined particles were then subjected to a C1 refinement with a shaped Fab-trimer mask and local angular searches only, followed by a third round of 3D classification without alignment using a shaped Fab-trimer mask. 3D classes that displayed distinct orientations of the Fab relative to the Env protomer were individually selected, and/or classes deemed sufficiently similar were combined, and their particles subjected to individual refinements with local angular searches only and post-processing.

### Cryo-EM

Purified HIV-1 Env stabilized CH505 TF chimeric SOSIP Env trimer (CH505 TF chTrimer) preparations were diluted to a final concentration of about 1 mg/mL in 2 mM Tris pH 8.0, 200 mM NaCl and 0.02% sodium azide, were mixed with 6-fold molar excess of DH1285 Fab and incubated for 2 hours at room temperature. 2.5 µL of protein was deposited on a Quantifoil 1.2/1.3 holey carbon grid that had been glow discharged for 30 seconds in a PELCO easiGlow Glow Discharge Cleaning System. After a 30 second incubation in > 95% humidity, excess protein was blotted away for 2.5 seconds before the grid was plunge frozen into liquid ethane using a Leica EM GP2 plunge freezer (Leica Microsystems). Cryo-EM data were collected on a FEI Titan Krios microscope (Thermo Fisher Scientific) operated at 300 kV. Data were acquired with a Gatan K3 detector operated in counting mode. Data processing was performed within cryoSPARC^52^ including particle picking, multiple rounds of 2D classification, *ab initio* reconstruction, heterogeneous and homogeneous map refinements, local refinement, and non-uniform map refinements. ChimeraX ^53^, Coot^54^, Isolde^55^ and Phenix^56^ were used for model-building and refinement.

### Epitope-Specific Single B Cell Sorting

Peripheral blood B cell sorting was performed as described previously^57,58^. Briefly, cryopreserved PBMC were stained with viability dye, CD14, CD16, CD20, CD3, CD27, IgD, fluorophore-labeled Man_5_GlcNAc_2_-enriched CH505.M5.G458Y, and fluorophore-labeled Man_5_GlcNAc_2_-enriched CH505.M5.G458Y.N280D Env Trimer. Env trimers with C-terminal avi-tags (Avidity) were biotinylated and conjugated to streptavidin labeled with different fluorochormes. Live, IgD-negative single B cells that bound to CH505.M5.G458Y but not CH505.M5.G458Y.N280D were sorted into cell lysis buffer and 5X first-strand synthesis buffer in individual wells of a 96-well PCR plate. Plates were frozen on dry ice and ethanol immediately and stored at −80 °C until reverse transcription of RNA.

### Rhesus Immunoglobulin RT-PCR

Immunoglobulin genes were amplified as previously described^57,58^. Immunoglobulin genes from a single B cell were reverse transcribed with Superscript III (ThermoFisher) and constant region-specific reverse primers. Five microliters of complementary DNA were used for two rounds of nested PCR. PCR amplicons were purified and sequenced with 4 _μ_M of forward and reverse primers. Contigs of the forward and reverse antibody sequences were made by the Duke automated sequence analysis pipeline (Duke ASAP). Immunogenetics of rhesus macaque immunoglobulin genes were determined with the macaque heavy and light chain reference library in Cloanalyst, where IGHV1-h is the closest v-gene segment based on sequence identity to human IGHV1-46. Rhesus IGHV gene segments were subsequently annotated using the macaque sequence database in KIMDB where genes denoted IGHV1-h by Cloanalyst were called as IGHV1-105*01^39,59^. Recombination summaries of the immunogenetics of identified antibodies were partitioned into clones and an unmutated common ancestor (UCA) was inferred from select clones with the infer UCA function in Cloanalyst. A phylogenetic tree for the DH1285 lineage was generated using the heavy chain sequences since light chain gene clonality has a certain degree of uncertainty due to the single V and J junction used to determine clonal relatedness. Each antibody with interpretable sequencing was expressed via a linear DNA cassette in Expi293F cells (Thermo Fisher Scientific, Cat No. A14527). Cell culture media was tested for binding to HIV-1 envelope. Antibodies with binding or immunogenetics of interest were synthesized and cloned into gamma, kappa, or lambda expression vectors (GenScript). Plasmids were prepared for transient transfection of Expi293F cells using the MidiPrep plasmid plus kit (Qiagen). Antibodies were produced in Expi293F cells (Thermo Fisher Scientific, Cat No. A14527).

### Next generation sequencing of antibody genes

Illumina MiSeq sequencing of antibody heavy chain VDJ and VK sequences was performed on peripheral B cells from week 32 as previously described^49^. Total RNA was isolated from PBMCs and two independent cDNA samples were generated for library construction. We expect sequences error due to NGS sample preparation to be less than four base pairs (<1%) based on previous experiments. Thus, inference of the unmutated common ancestor and intermediate antibodies was performed with sequences observed in sequencing runs of both cDNA samples. V, D, and J gene segment inference, clonal relatedness testing and reconstruction of clonal lineage trees were performed using the Cloanalyst software package^59^.

### Biolayer interferometry (BLI)

Biolayer interferometry was performed as previously described^60^. BLI ligand titration and binding kinetics assays were performed on an Octet Red96e system (Sartorius) at 30°C with an orbital shake speed of 1000 rpm. Assays were performed in flat bottom 96-well plates with assays (Greiner) using 0.22μm-filtered Phosphate buffered saline supplemented with 0.05% Tween 20 and 0.1% bovine serum albumin (PBS-T-BSA). For the ligand titration experiment, two-fold serial dilutions of the biotinylated avi-tagged CH505 TF chSOSIPv4.1 and Man_5_GlcNAc_2_-enriched CH505 M5.G458Y chSOSIPv4.1 Env trimers were immobilized on hydrated Streptavidin (SA) Tips (Octet^®^ Sartorius) for 120 seconds. Env immobilization concentration started at 10 μg/mL. After a 60 second wash and 180 second Baseline step in PBS-T-BSA, the tips were incubated with 1000nM or 500 nM of IgG or Fab for 300 seconds. The optimal Env concentrations for affinity studies were analyzed from the sensorgram traces and binding responses using Data Analysis HT 12.0 software (Forte Bio). Ligand loading conditions where ligand loading was linear and Fab association reached approximately 0.4 nm were selected. Binding kinetics experiment were performed with optimal Env concentrations. Env was immobilized on hydrated SA biosensor tips and incubated with two-fold serial dilutions of antibody IgG or Fabs for 300 seconds followed by a 600-second-long dissociation step in PBS-T-BSA. The binding response sensorgram curves were globally fitted using a 1:1 Binding Model and the rate constants k_a_, k_d_ and K_d_ were calculated from 4 or more curves using Data Analysis HT 12.0 software (Forte Bio).

### Indirect ELISA Using Env Trimers

Corning 384-well plates were coated overnight at 4 °C with 15 µL of either streptavidin (for biotinylated proteins) or the base-binding RM19R, expressed with a human IgG constant region diluted to 2 µg/mL in 0.1M NaCO_3_. Plates were washed with PBS-T (PBS + 0.05% Tween-20), then all sample wells were blocked for 1 hour at room temperature (all subsequent steps) with 40 µL blocking buffer (PBS + 15% v/v goat serum + 4% w/w whey protein + 0.05% v/v Tween-20). Plates were again washed, then incubated with 15 µL/well Env diluted to 2 µg/mL in blocking buffer for 1 hour. Plates were washed again, then incubated with 10 µL of a serial dilution of rhesus antibodies or serum beginning at a concentration of 100 µg/mL or 1:30, respectively in blocking buffer for 1.5 hours. Samples were again washed, then incubated with anti-rhesus IgG conjugated to HRP (Southern Biotech Cat. No.: 4700-05) diluted in blocking buffer for 1 hour. Plates were again washed, then developed with 20 µL/well of a tetramethylbenzidine peroxidase substrate (SeraCare) for 15 minutes, then quenched with equal volume 1% HCl. Absorbance was measured at 450 nm on a SpectraMax 340 PC.

### Indirect ELISA Using Env gp120

Corning 384-well plates were coated overnight at 4 °C with 15 µL of gp120 diluted to 2 µg/mL in 0.1M NaCO_3_. Plates were washed with PBS-T, then all sample wells were blocked for 1 hour at room temperature (all subsequent steps) with 40 µL blocking buffer. Plates were washed again, then incubated with 10 µL of a serial dilution of rhesus antibodies or serum beginning at a concentration of 100 µg/mL or 1:30, respectively in blocking buffer for 1.5 hours. Samples were again washed, then incubated with anti-rhesus IgG conjugated to HRP (Southern Biotech Cat. No.: 4700-05) diluted in blocking buffer for 1 hour. Plates were again washed, then developed with 20 µL/well of a tetramethylbenzidine peroxidase substrate (SeraCare) for 15 minutes, then quenched with equal volume 1% HCl. Absorbance was measured at 450 nm on a SpectraMax 340 PC.

### Competition ELISAs

Corning 384-well plates were coated overnight at 4 °C with 15 µL streptavidin diluted to 2 µg/mL in 0.1M NaCO_3_. Plates were washed with PBS-T, then all sample wells were blocked for 1 hour at room temperature (all subsequent steps) with 40 µL blocking buffer (PBS + 15% v/v goat serum + 4% w/w whey protein + 0.05% v/v Tween-20). Plates were again washed, then incubated with 15 µL/well Env diluted to 2 µg/mL in blocking buffer for 1 hour. Plates were washed again, then incubated with 10 µL/well 2G12 or CH235.12 (human IgG) diluted to 50 µg/mL in blocking buffer. Plates were washed again, then incubated with 10 µL of a serial dilution of rhesus antibodies or serum beginning at a concentration of 100 µg/mL or 1:30, respectively in blocking buffer for 1.5 hours. Samples were again washed, then incubated with anti-rhesus IgG conjugated to HRP (Southern Biotech Cat. No.: 4700-05) diluted in blocking buffer for 1 hour. Plates were again washed, then developed with 20 µL/well of a tetramethylbenzidine peroxidase substrate (SeraCare) for 15 minutes, then quenched with equal volume 1% HCl. Absorbance was measured at 450 nm on a SpectraMax 340 PC.

## Acknowledgments

We thank Victoria Gee-Lai, Margaret Deyton, Advaiti Khanore, Giovanna Hernandez, and Aja Sanzone for technical assistance. We thank Elizabeth Donahue for program management and assistance with manuscript preparation. Illumina NGS was performed by the DHVI Viral Genetics Analysis Core Facility. Flow cytometry was performed by the DHVI Flow Cytometry Core Facility.

## Funding

This study was funded by the National Institutes of Health, National Institute of Allergy and Infectious Diseases, Division of AIDS grant UM1AI144371 for the Duke Consortia for HIV/AIDS Vaccine Development (BFH), U54 AI170752 (PA), and R01 AI145687 (PA).

## Author contributions

KOS, DW and BFH conceived the macaque immunogenicity study. ML, SS, and LS administered the immunogenicity study and processed samples from the macaque study. JC, RP, MB and NH characterized antibody binding and analyzed results. XL isolated and sequenced the antibodies. JC, NJ, MK, RH, and JB performed protein production for antibodies and envelopes. RE and KatM performed negative stain electron microscopy experiments. VS, KJ, KarM, BT, and PA determined cryo-electron microscopy structures. YC, BH, WBW performed next-generation sequencing of antibody genes. KW and SV analyzed antibody sequences and performed bioinformatic inferences of clonality. KA, and SMA performed binding kinetics assays with surface plasmon resonance. JB and KOS performed binding kinetic assays with biolayer interferometry. YT and CB provided lipid nanoparticle adjuvant. CJ, AE, and DCM generated and analyzed serum and monoclonal antibody neutralization results. MAM and DWC performed fluorescence-activated cell sorting and flow cytometry phenotyping experiments. BFH, KOS, PA, BT, RE, and JC wrote the first draft of the manuscript, which was edited by all coauthors. KOS, JC, and BFH reviewed all study data. BFH provided funding for the study.

## Competing interests

YT and CB are employees of Acuitas Therapeutics. Acuitas Therapeutics had no role in the execution of the study, data collection, or data interpretation. KOS, DCM, RH, PA, and BFH have patents concerning the envelope immunogens used in this study. All remaining authors declare no competing interests.

## Data and materials availability

All data are available in the main text or the supplementary materials. Cryo-EM data sets have been deposited in the Protein database and electron microscopy database under accessioning numbers EMD-27621 and EMD-27622. All unique reagents generated in this study are available from the lead contact with a completed materials transfer agreement between the donor and recipient institutions.

## Lead contact

Further information and requests for resources and reagents should be directed to and will be fulfilled by the lead contact, Kevin Saunders.

