## Supplemental Figures and Tables for "Vaccine induction of CD4-mimicking broadly neutralizing antibody precursors in macaques"

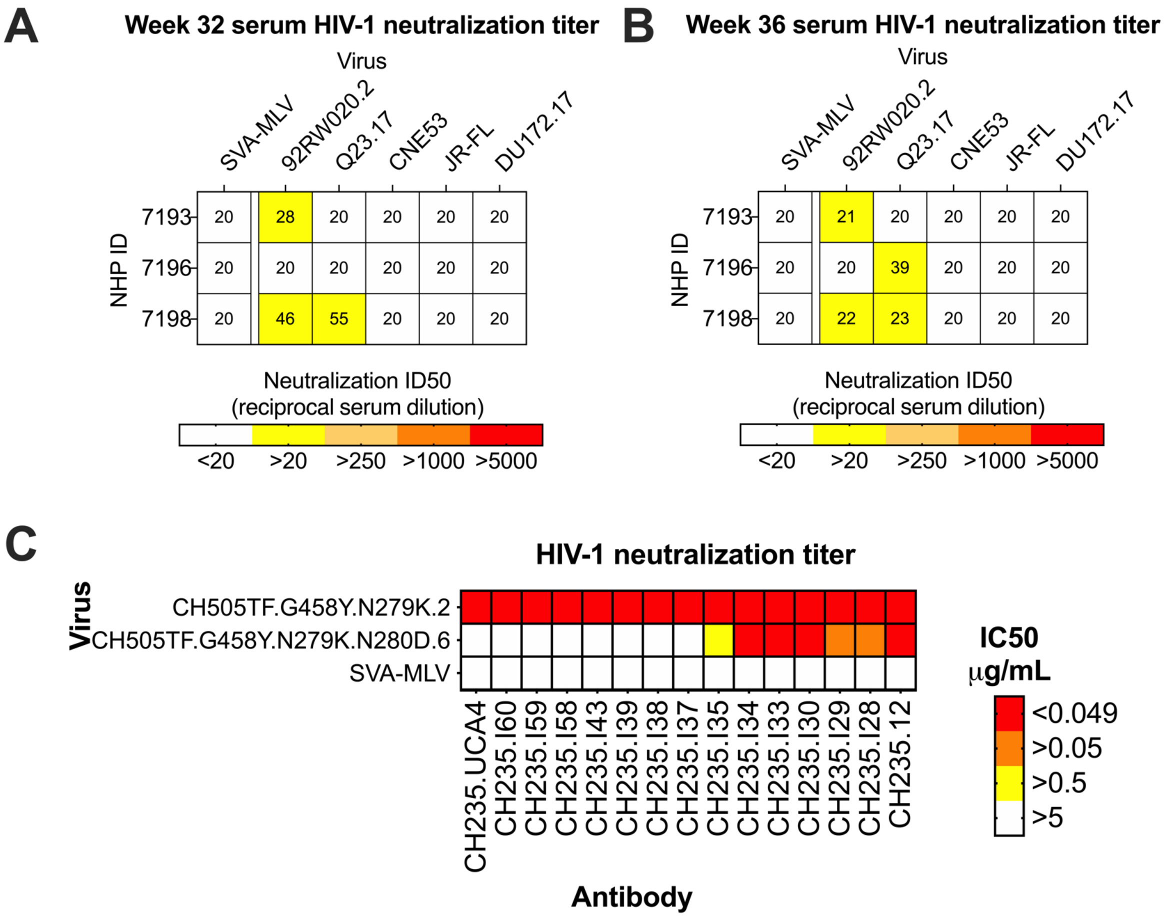


**Figure S1. CH235 lineage and vaccinated macaque sera neutralization characteristics. A-B)** Vaccinated macaque serum heterologous neutralization titers. Heatmap of vaccinated macaque **A)** week 32 and **B)** week 36 serum neutralization against heterologous envelope pseudotyped viruses, expressed as reciprocal serum dilution required to neutralize 50% of virus replication. **C)** CH235 lineage dependence on Asparagine at position 280 (N280) in the CD4bs. CH235 lineage antibodies develop neutralization against the escape mutant N280D in CH505.M5.G458Y pseudotyped virus by intermediate I35. Values represent neutralization titers expressed as inhibitory concentration required to neutralize 50% of virus replication. Related to Figure 1.

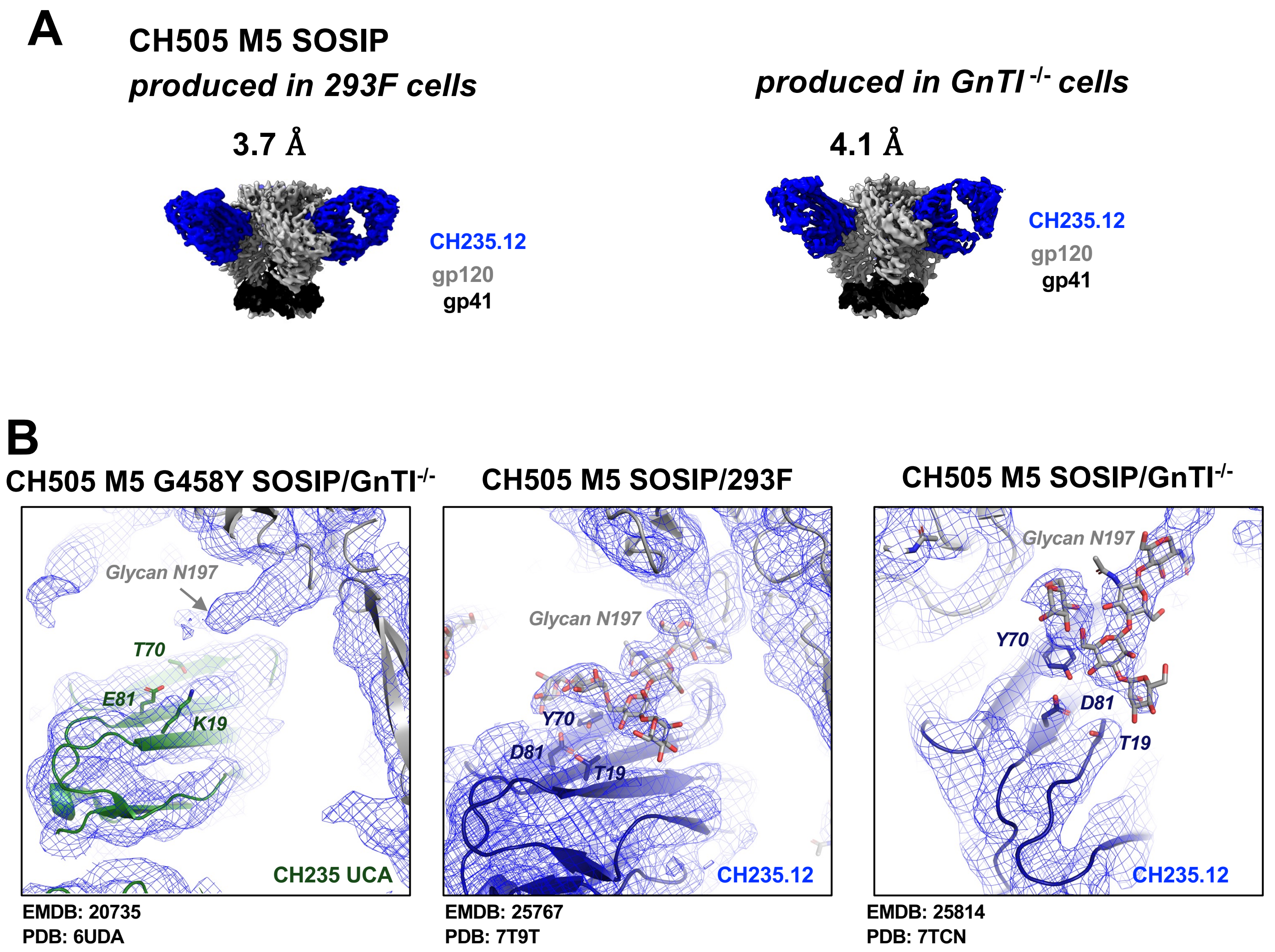


**Figure S2. Cryo-EM structures of CH235.12 bound to CH505 SOSIP Env (A)** Cryo-EM reconstructions of CH235.12 Fab bound to CH505 TF SOSIP with the N279K(M5) mutation in Loop D, where the SOSIP was produced either in 293F cells (left) or in GnT1- cells (right). **(B)** Zoomed-in view of the region around glycan N197 shown for (left) CH235 UCA bound to CH505 M5 G458Y SOSIP Env produced in GnT1- cells (EMDB: 20735; PDB: 6UDA) with heavy chain residues K19, T70 and E81 shown in sticks. The cryo-EM map density is shown as a blue mesh. The density for the disordered glycan N197 is indicated with an arrow, (middle) CH235.12 bound to CH505TF M5 SOSIP Env produced in 293F cells (EMDB: 25767; PDB: 7T9T) with heavy chain residues T19, Y70 and D81 and Env glycan 197 shown in sticks, (right) CH235.12 bound to CH505TF M5 SOSIP Env produced in GnT1- cells (EMDB: 25814; PDB: 7TCN) with heavy chain residues T19, Y70 and D81 and Env glycan 197 shown in sticks. Related to Figure 1.

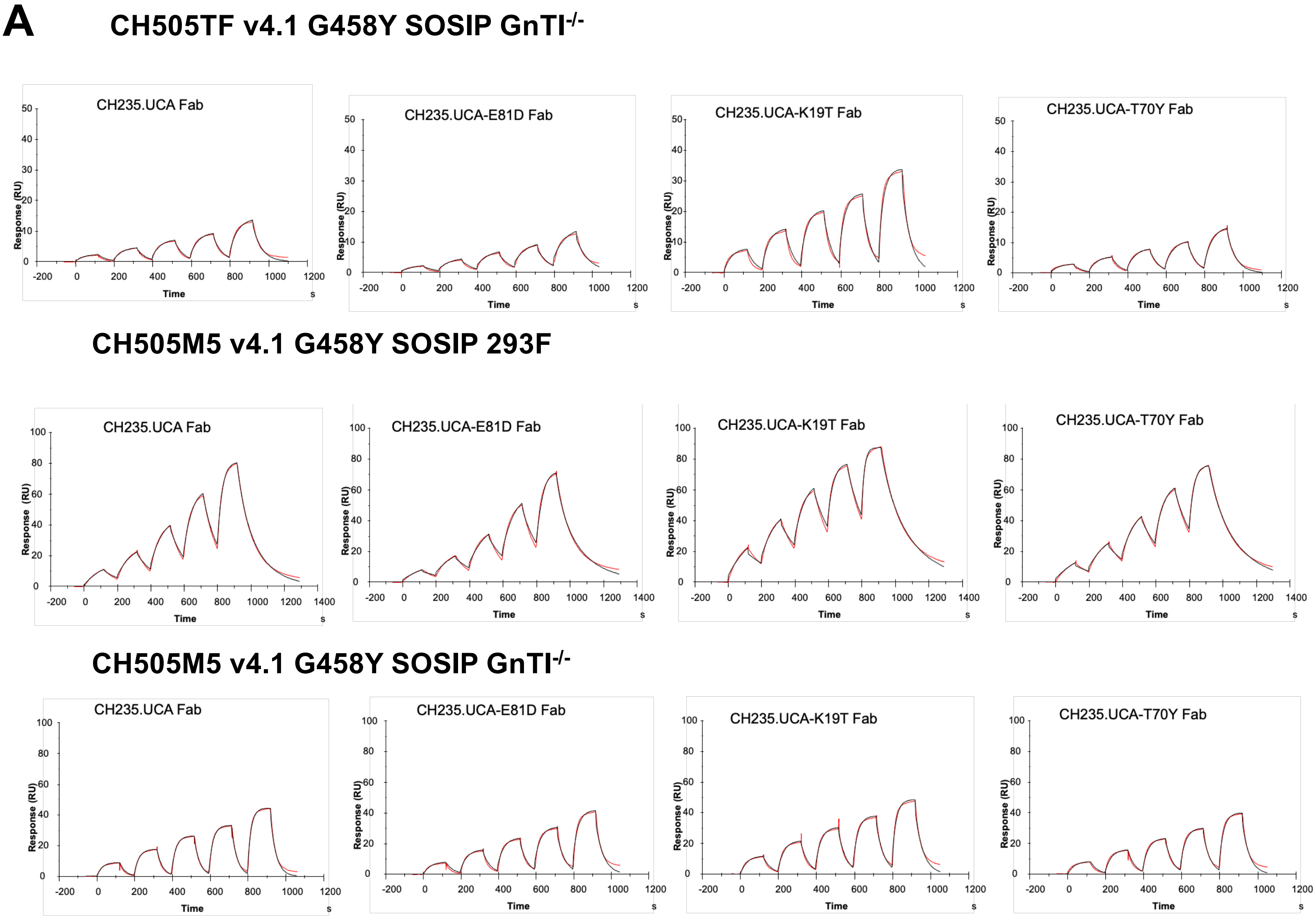


**B**

| | CH505TF v4.1 G458Y SOSIP GnTI$^{-/-}$ | | | CH505M5 v4.1 G458Y SOSIP GnTI$^{-/-}$ | | | CH505M5 v4.1 SOSIP 293F | | |
|---|---|---|---|---|---|---|---|---|---|
| Fab | ka ($M^{-1}s^{-1}$) | kd ($s^{-1}$) | KD (μM) | ka ($M^{-1}s^{-1}$) | kd ($s^{-1}$) | KD (μM) | ka ($M^{-1}s^{-1}$) | kd ($s^{-1}$) | KD (μM) |
| CH235 UCA | 936.1 | 0.02195 | 23.45 | 4.865E+4 | 0.01495 | 0.307 | 7173 | 0.03360 | 4.7 |
| CH235 UCA E81D | 287.4 | 0.01511 | 52.59 | 7.423E+4 | 0.02611 | 0.352 | 4788 | 0.02447 | 5.1 |
| CH235 UCA K19T | 6052 | 0.02282 | 3.8 | 8.048E+4 | 0.008268 | 0.103 | 8184 | 0.02409 | 2.9 |
| CH235 UCA T70Y | 877.6 | 0.02667 | 24.74 | 5.060E+4 | 0.009314 | 0.184 | 5615 | 0.02667 | 4.8 |

**Figure S3. Binding analysis to probe the effect of heavy chain mutations on Env binding (A)** Binding of CH235 UCA mutants to CH505 Env measured using surface plasmon resonance (SPR). The black lines show raw data, and the red lines are global fits of the data to a 1:1 Langmuir binding model. **(B)** Affinity and kinetics of CH235 UCA mutants binding to CH505 Env.

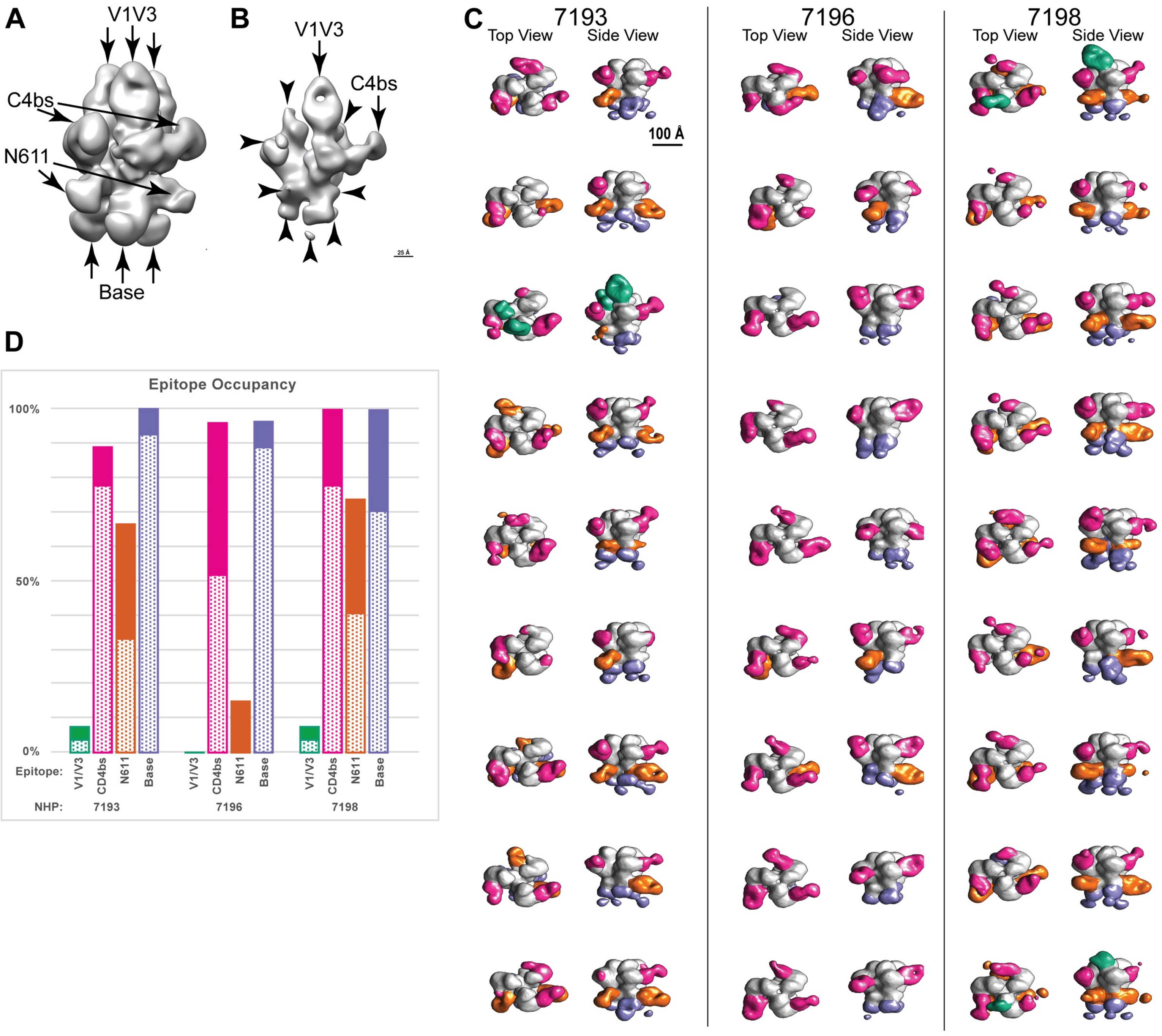


**Figure S4. Structural repertoire of envelope-specific antibodies in serum from vaccinated macaques.** (**A-B**) 3D reconstruction of negative stain electron microscopy (NSEM) images of serum-derived Fabs bound to CH505 M5.G458Y Env trimers. (**A**) Low and (**B**) high contour views of envelope in complex with multiple Fabs. Arrows indicate Fabs bound to envelope. Fabs bound to the CD4bs and V1V3 are denoted. (**C**) Top and side views of 3D reconstructions of serum Fabs in complex with CH505 M5.G458Y Env trimers. Fabs are colored based on specificity (V1/V3, green; CD4bs, magenta; N611, orange; base, purple). The ID number of the vaccinated macaque from which the serum Fabs were purified is shown at the top of each column. Each row shows a different structural class of Env trimer and Fab complex. **(D)** Percentage of Env:Fab particles containing each Fab specificity. Related to Figure 1.

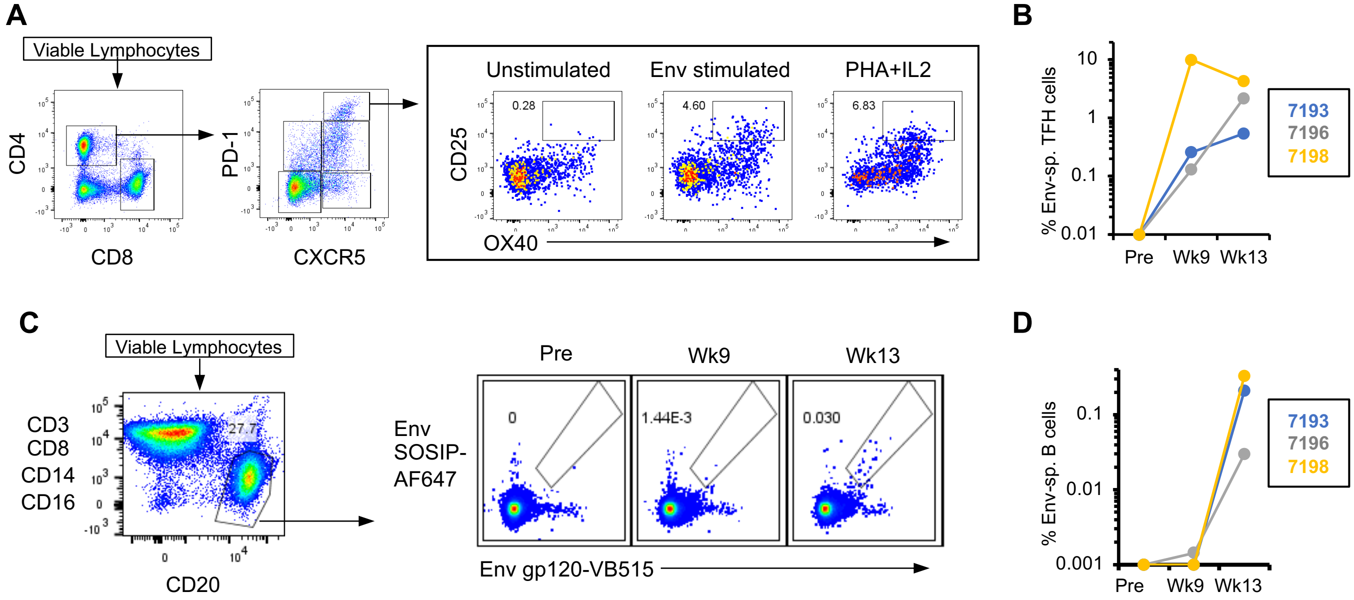


**Figure S5. Vaccination elicits Env-specific B lymphocytes and T follicular helper (TFH) cells in macaques. A)** Representative flow cytometry showing the gating strategy for identifying envelope-specific TFH CD4+ T cells in lymph nodes. Cells were stimulated with Env peptides covering the whole CH505 TF Env gp140 trimer or with PHA and IL-2 as a positive control. **B)** Quantification of Env-specific CD4+ TFH cells at 9 weeks (post 2 immunizations) and 13 weeks (post 3 immunizations). **C)** Representative flow cytometry gating strategy for the identification of Env-specific B cells in lymph nodes. **D)** Quantification of Env-specific B cells at 9 weeks (post 2 immunizations) and 13 weeks (post 3 immunizations) in lymph nodes. Related to Figure 2.

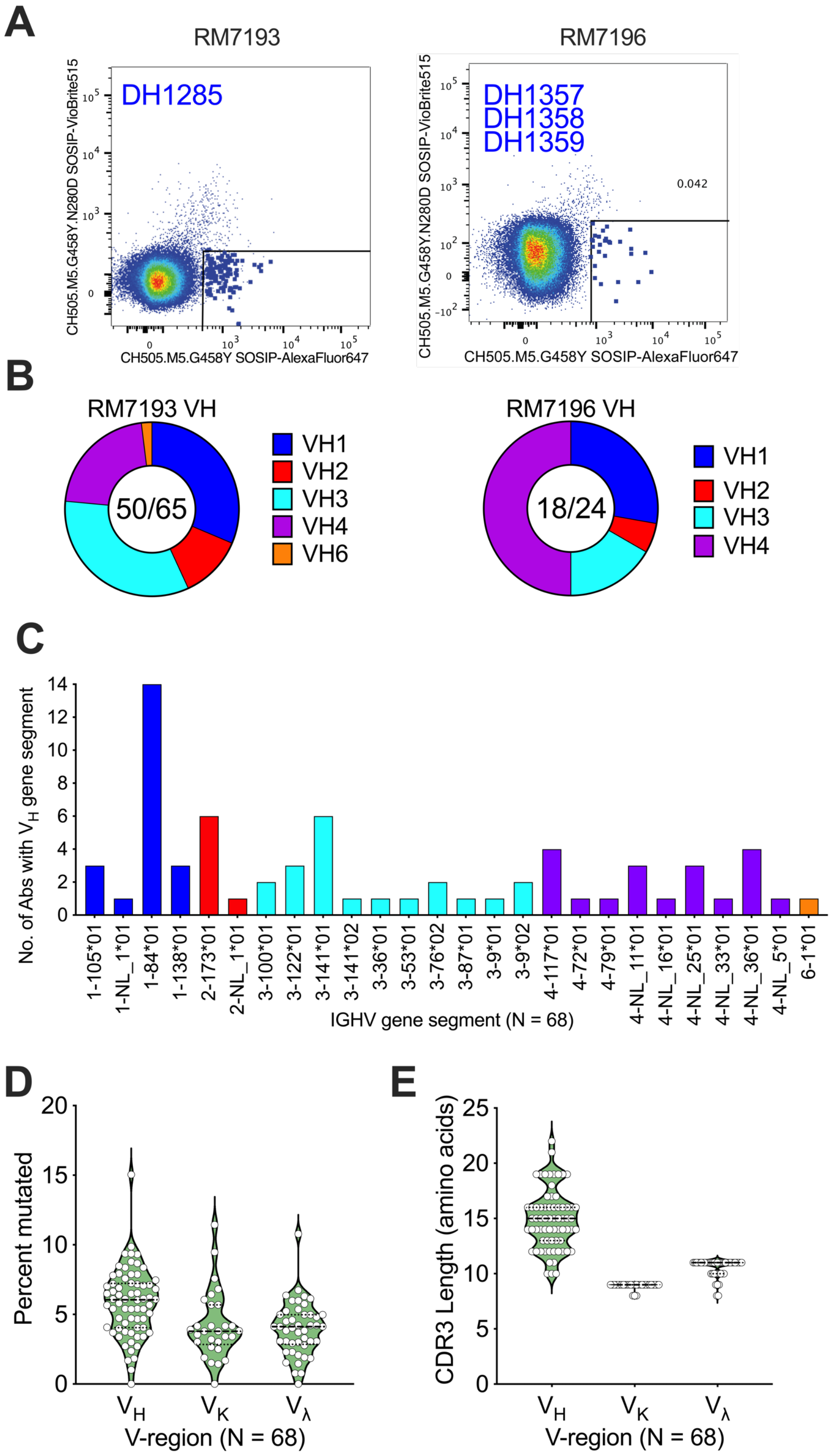


**Figure S6. Detailed isolation and immunogenetics of Env-specific monoclonal antibodies from animals 7193 and 7196. A)** Fluorescence activated cell sorting dot plot showing sorting of N280D-dependent B cells specific for CH505.M5.G458Y SOSIP trimer. CH235-like antibodies from each animal are listed. **B)** Identification of heavy chain variable gene segment family usage among isolated monoclonal antibodies. The number of unique B cell clones with interpretable full variable region sequence is shown as the first number inside the pie chart followed by the total number of B cell clones identified. **C)** Frequency of KIMDB heavy chain variable gene segment assignment among monoclonal antibodies. **D,E)** Violin plot of the distribution of heavy and light chain variable region (**D**) nucleotide mutation and (**E**) CDR3 lengths observed for antibodies from macaques 7193 and 7196. Related to Figure 2.

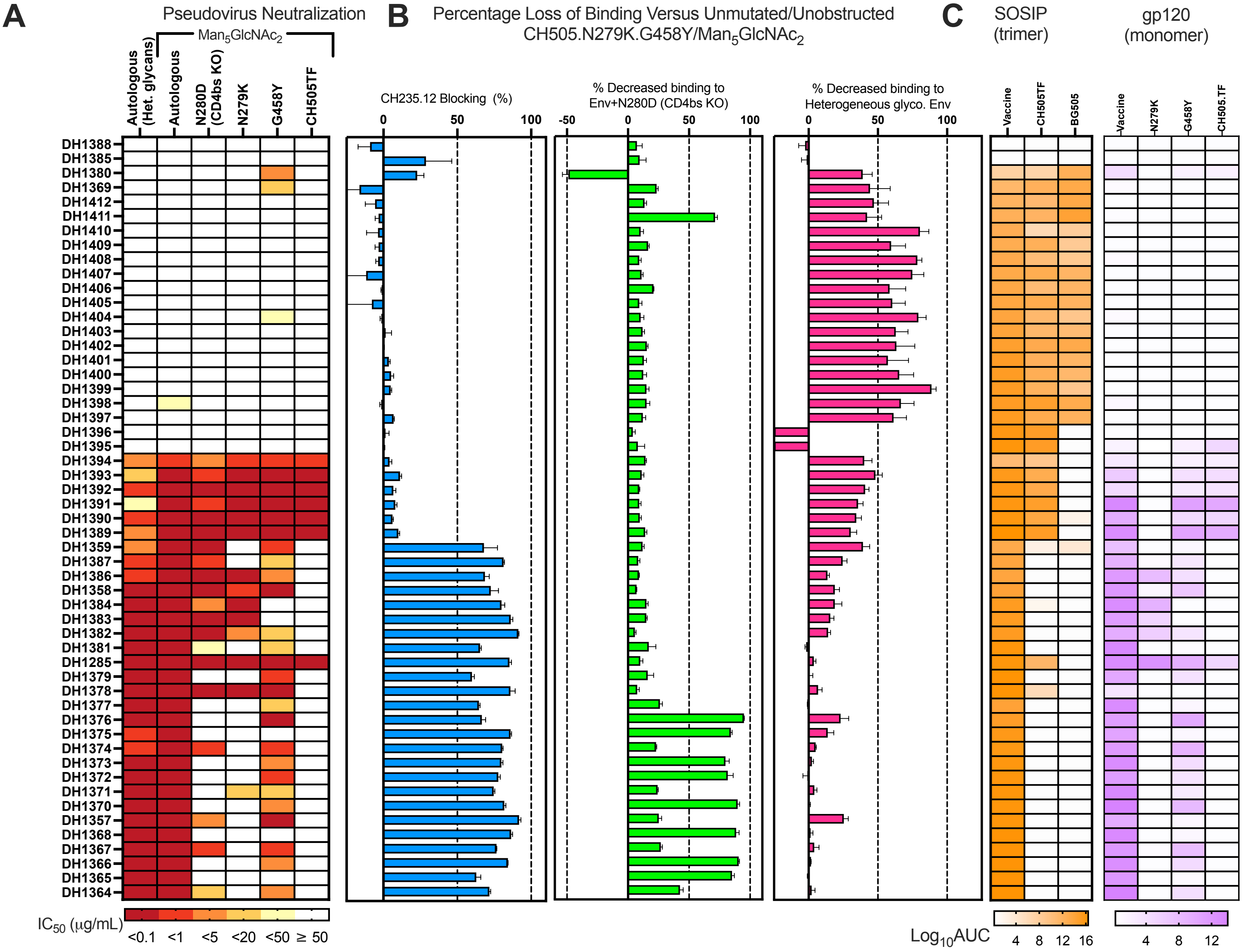


**Figure S7. Neutralization and binding characteristics of Env-specific monoclonal antibodies from vaccinated macaques. A)** Neutralization titer (as inhibitory concentration required for 50% neutralization) against different pseudotyped CH505 viruses. The autologous virus matches the M5.G458Y envelope used in the vaccine and was produced with heterogenous (Het.) glycans or $Man_5GlcNAc_2$-enriched. $Man_5GlcNAc_2$-enriched M5.G458Y with a CD4bs knockout (KO) mutation (N280D) was also tested. Neutralization titers are shown for $Man_5GlcNAc_2$-enriched TF with a N279K (M5), G458Y, or no substitutions. **B)** Percentage loss of monoclonal antibody binding in the presence of CH235.12 blocking (blue), the CD4bs resistance mutation N280D (green), or with heterogenous glycosylation (magenta). All assays use or compare binding against CH505.M5.G458Y/GnT1$^-$. Error bars represent the standard deviation of three replicates. **C)** Binding magnitude of monoclonal antibodies for $Man_5GlcNAc_2$-enriched CH505.M5.G458Y (autologous to the vaccine), CH505.TF without glycan enrichment, or BG505 without glycan enrichment SOSIP trimers (orange). Binding magnitude for monomeric gp120s of CH505 Env derivatives is shown on the right (purple). Binding magnitude is shown as area under the $\log_{10}$ transformed curve (logAUC). Related to Figure 2.

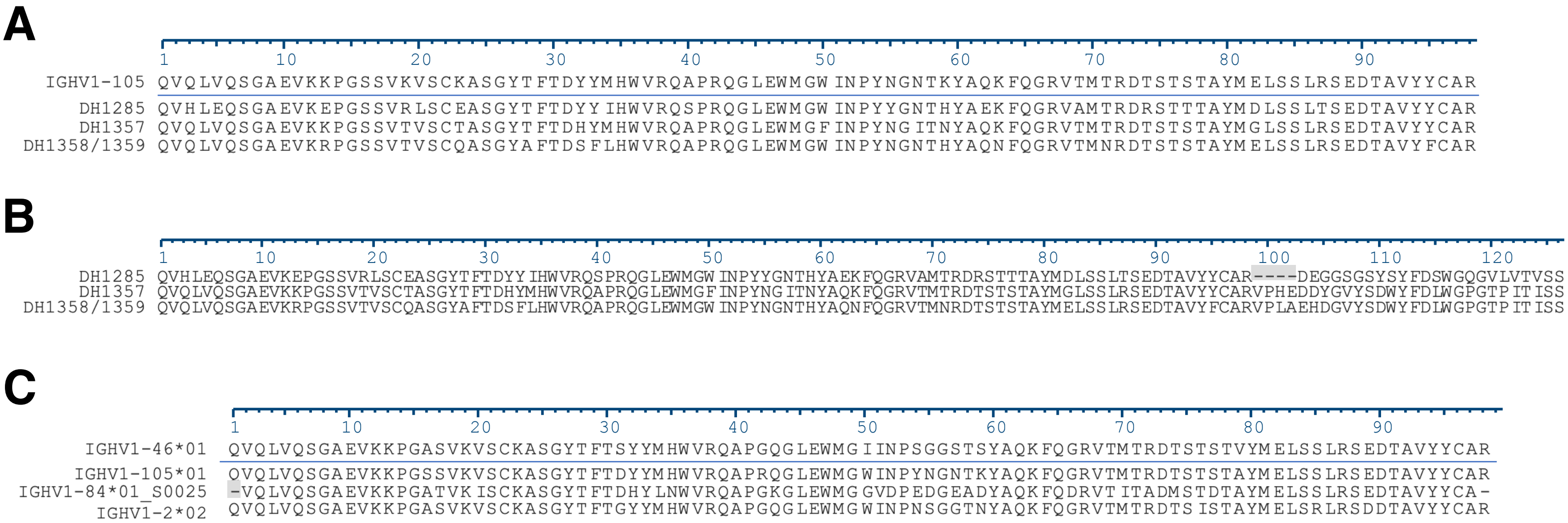


**Figure S8. Multisequence alignment of heavy chain variable regions of CD4bs antibodies from vaccinated rhesus macaques. A)** Comparison of vaccinated macaque heavy chain V gene segments and the inferred KIMDB germline reference gene segment IGHV1-105. DH1358 and DH1359 are composed of the same heavy chain variable region sequence. (**B**) Comparison of IGHV1-105-derived macaque CD4bs variable region amino acid sequences. Dashes indicate a gap in the CDR3 of DH1285 relative to DH1357 and DH1358. **C)** Comparison of human variable regions used by CD4 mimicking bnAbs (IGHV1-46*01 GenBank accession number X92343 and IGHV1-2*02 GenBank accession number X62106) and rhesus macaque variable regions observed from Env-specific B cells. IGHV1-84 was the most abundant allele observed among Env-specific B cells, but is only 78% identical to rhesus IGHV1-105*01 or human IGHV1-46*01 and 76% identical to human IGHV1-2*02. Rhesus IGHV1-105*01 and human IGHV1-46*01 were 91% identical to each other. Related to Figure 3.

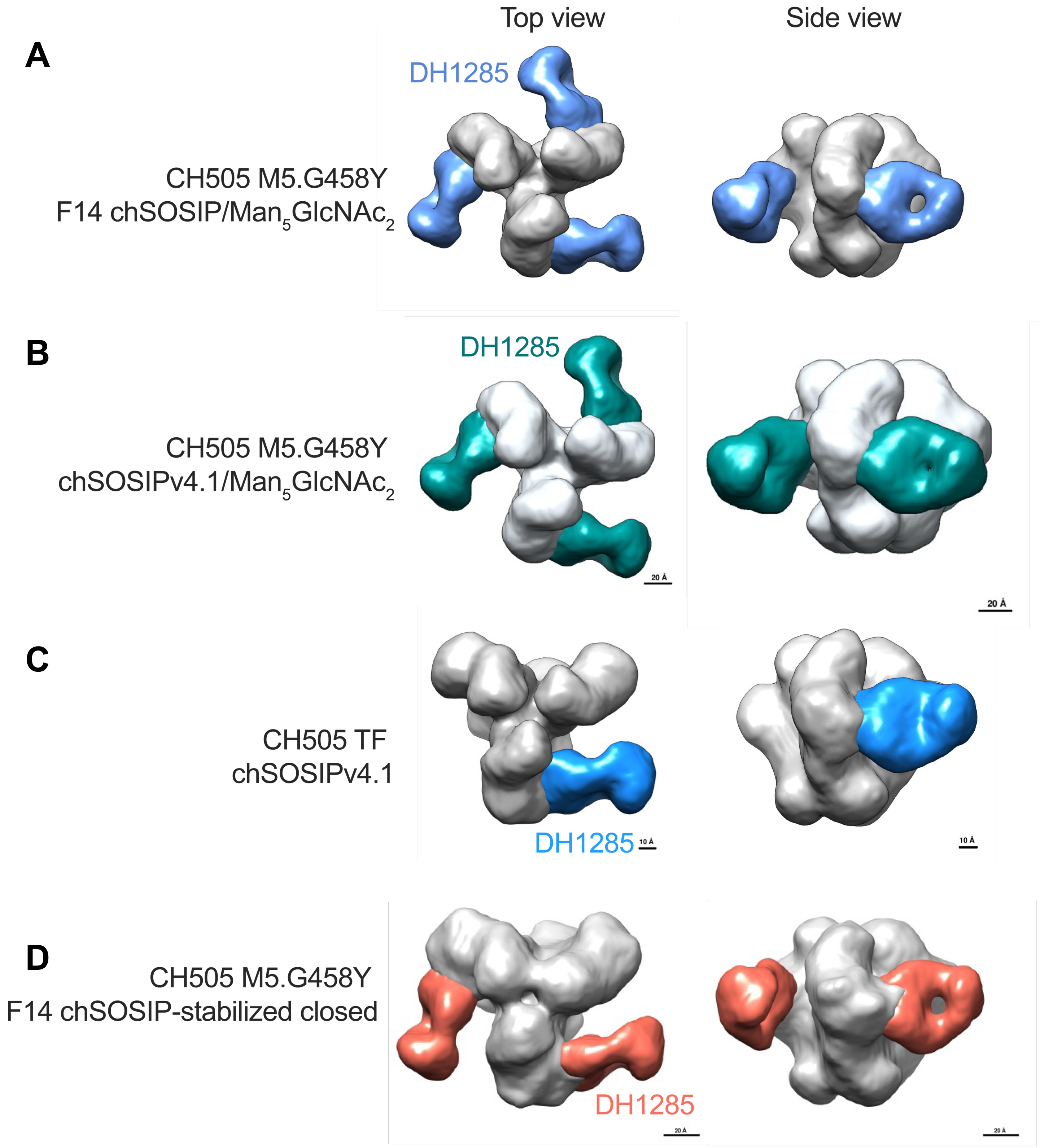


**Figure S9. DH1285 binds to partially open or closed conformations of envelope. A-C)** 3D reconstruction from NSEM images of DH1285 bound to partially open F14-stabilized CH505.M5.G458Y with $Man_5GlcNAc_2$ enrichment, SOSIPv4.1-stabilzed CH505.M5.G458Y with $Man_5GlcNAc_2$ enrichment, and CH505.TF SOSIPv4.1. **D)** Env trimer. DH1285 bound to closed Env is shown for a hyperstabilized M5.G458Y Env trimer. Env is shown in gray and DH1285 Fab is colored light blue, blue, teal, or red. Related to Figure 3.

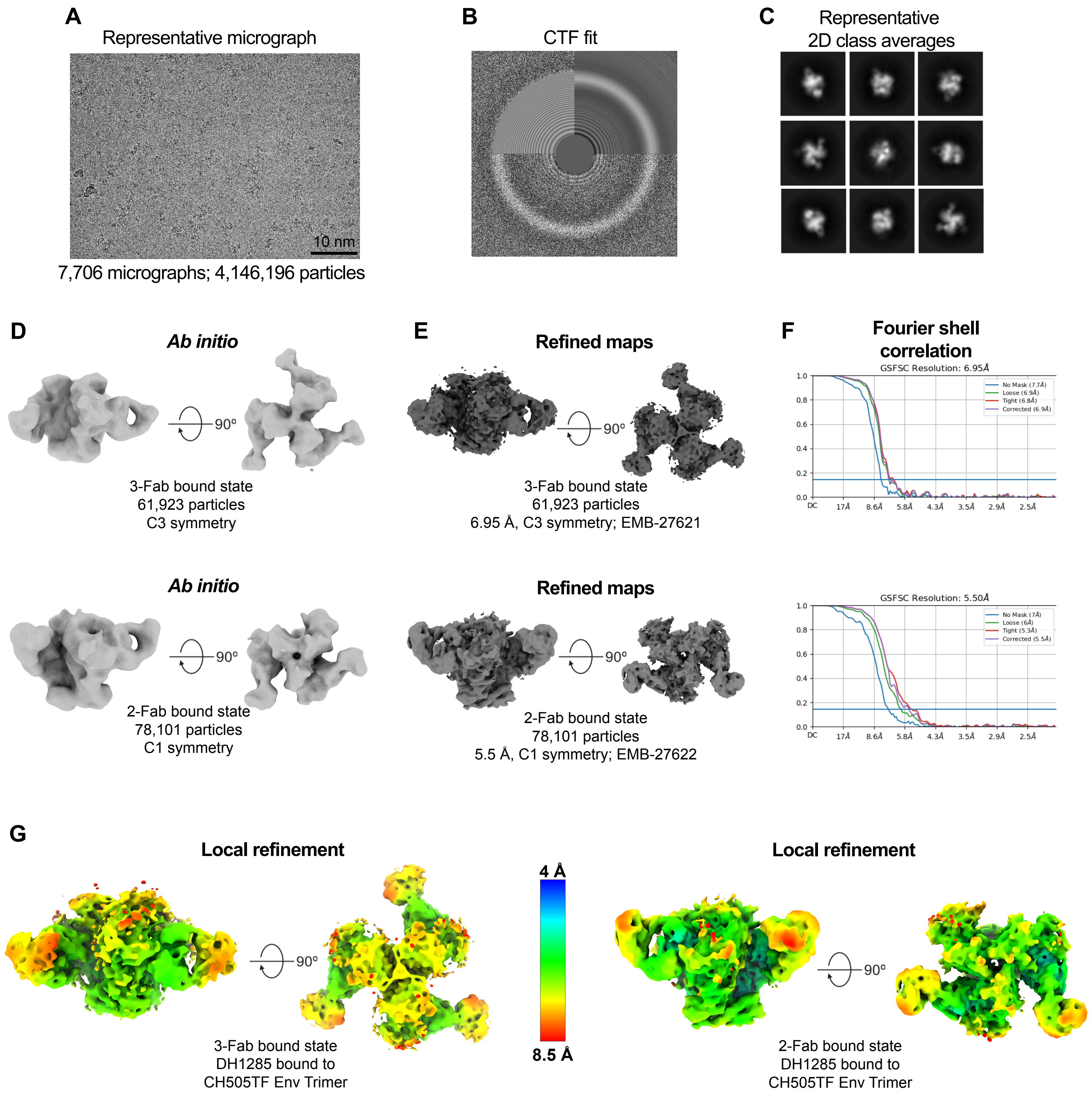


**Figure S10. Cryo-EM data processing for Antibody DH1285 in complex with HIV-1 Env trimer.** **(A)** Representative micrograph. **(B)** Representative CTF fit. **(C)** Representative 2D class averages from cryo-EM dataset. Box size = 345.6 Å. **(D)** Ab initio maps of 3-Fab and 2-Fab bound states. **(E)** Refined maps for corresponding states. **(F)** Fourier Shell Correlation (FSC) curves of the 3D reconstruction shown in E with horizontal blue line indicating FSC 0.143. **(G)** Refined map shown in E colored by local resolution. Related to Figure 4.

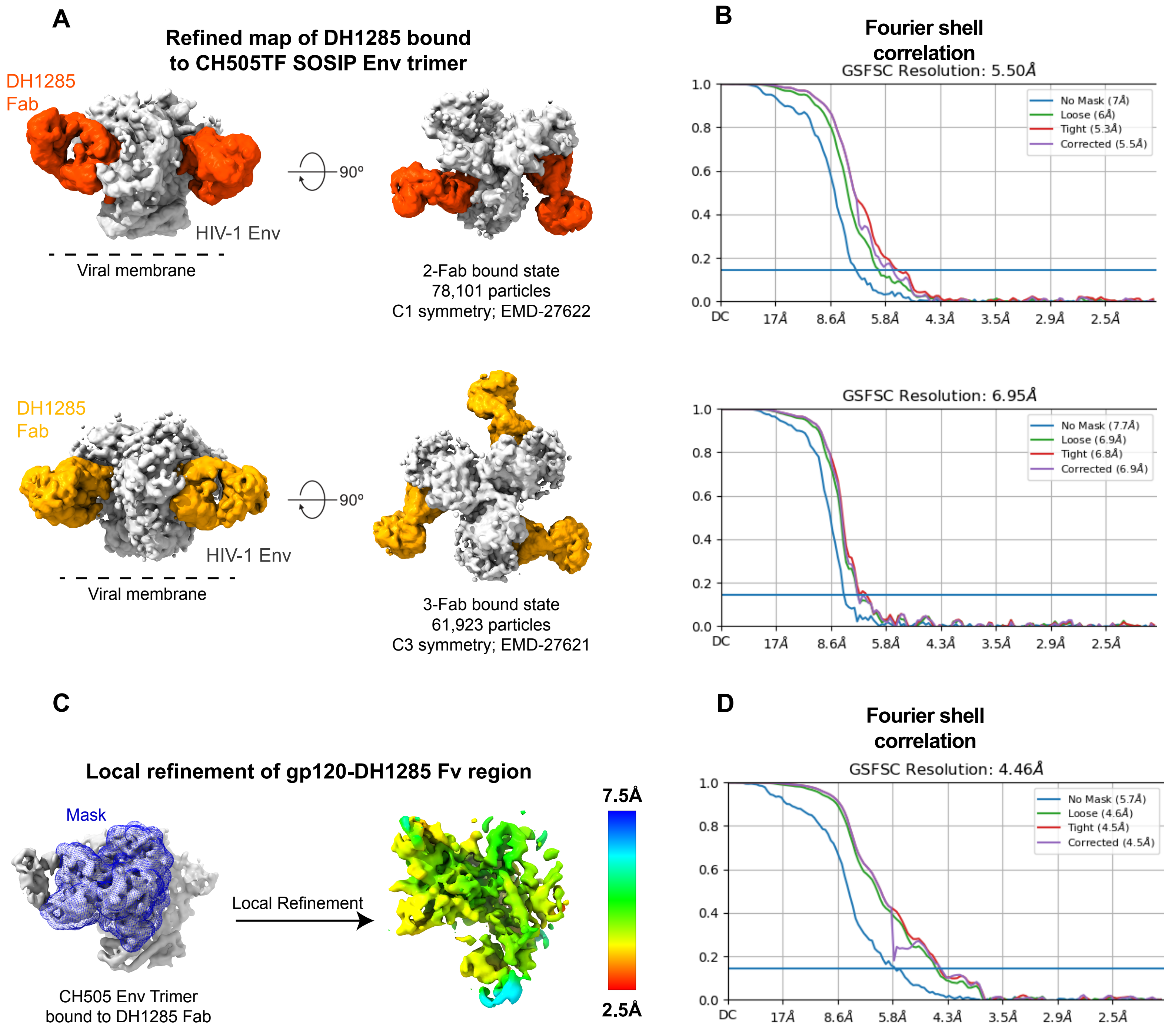


**Figure S11. Cryo-EM data processing for Antibody DH1285 in complex with HIV-1 Env trimer local refinement. (A)** Cryo-EM reconstructions of DH1285 Fab bound to HIV-1 Env trimer. 2-Fab bound state colored by segmented refined map, with Fab colored orange. 3-Fab bound state with Fab colored orange-yellow. **(B)** Corresponding Fourier Shell Correlation (FSC) curves of the 3D reconstruction shown in A with horizontal blue line indicating FSC 0.143. **(C)** Local refinement of gp120-DH1285 Fv region. (Left) mask is shown in blue mesh and made from EMD-27622 binding interface. (Right) local refinement map EMD-XXXX colored by local resolution ranging from 2.5 to 7.5 Å. **(D)** Fourier Shell Correlation (FSC) curve of local refinement shown in C with horizontal blue line indicating FSC 0.143. Related to Figure 4.

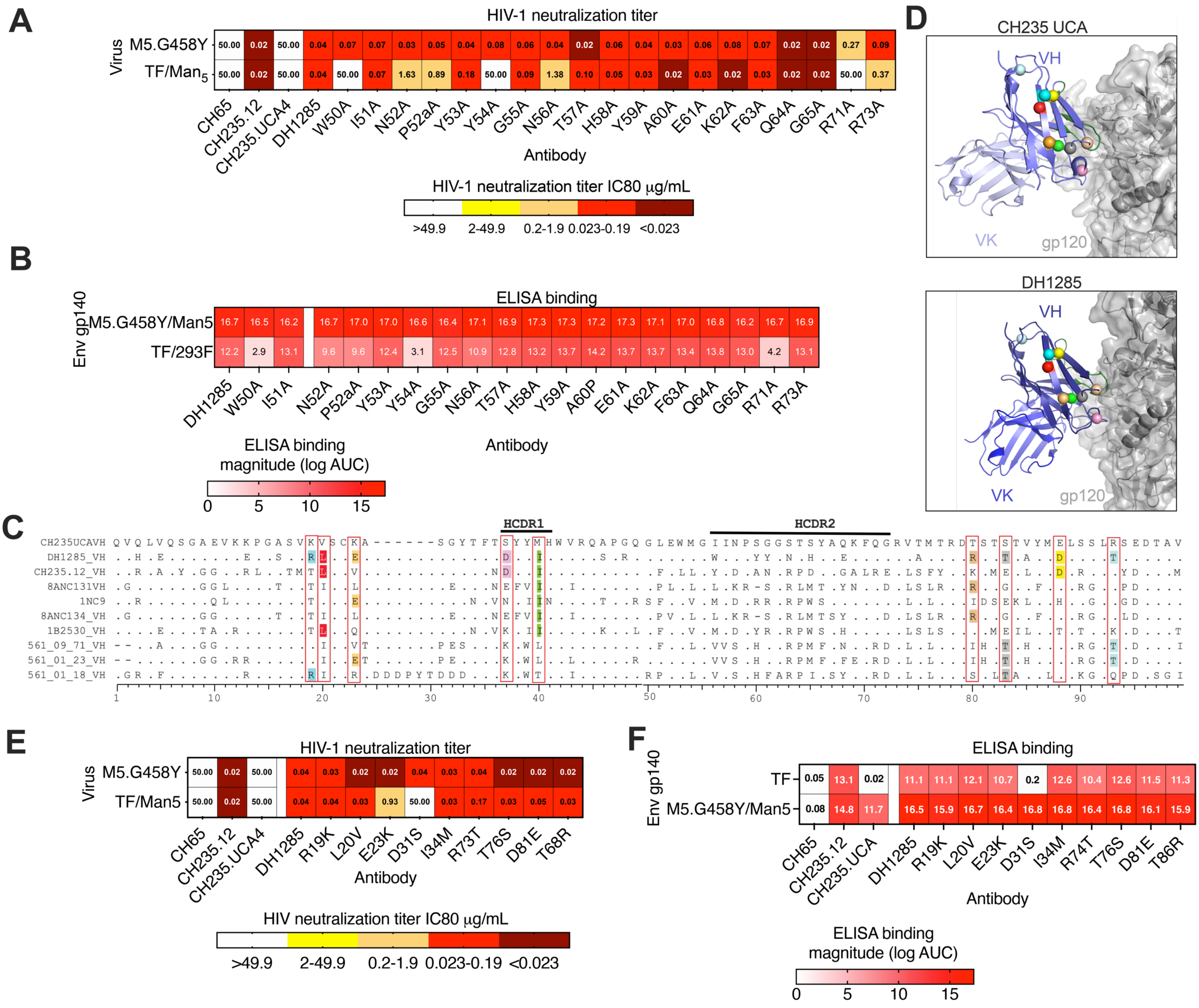


**Figure S12. Identification of the amino acids within the DH1285 paratope that are required for neutralization and binding to Env trimer. A)** CH505.TF and $Man_5GlcNAc_2$ enriched CH505.M5.G458Y pseudotyped virus neutralization by HCDR2 and framework region 3 DH1285 mutants. Titers are shown as antibody concentration required to inhibit 80% of virus replication. **B)** Binding magnitude of DH1285 mutant antibodies shown (A) for CH505.TF and $Man_5GlcNAc_2$ enriched CH505.M5.G458Y gp140 Env trimers by ELISA Binding magnitude is shown as log AUC. **C)** Identification of conserved amino acid residues of DH1285 compared to other $V_H$1-46 human bnAbs. Boxes indicate amino acid positions where the DH1285 amino acid is identical to one or more VH1-46 bnAbs. **D)** The structures of Env gp120 (gray) in complex with CH235 UCA (top) or DH1285 (bottom) show the location of VH1-46 bnAb class amino acids observed in DH1285. Amino acids conserved between VH1-46 bnAbs and DH1285 are shown as spheres colored as in **C**. **E)** Neutralization activity and **(F)** binding reactivity of DH1285 antibodies with substitutions at amino acids position conserved in VH1-46 bnAbs. DH1285 amino acids were substituted for the CH235 UCA amino acid. Related to Figure 5.

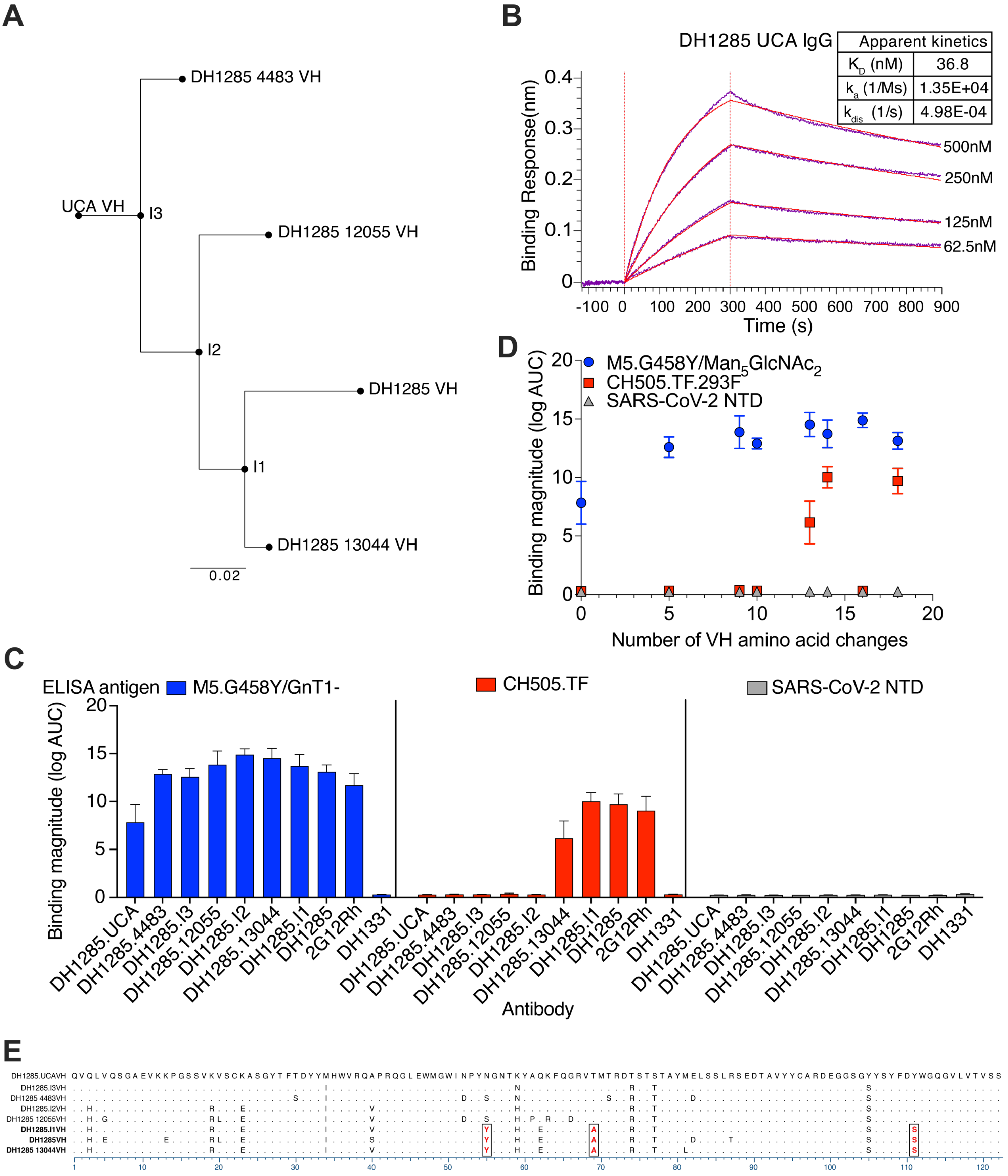


**Figure S13. DH1285 lineage Env binding activity. A)** Cloanalyst **i**nferred DH1285 lineage tree including members identified from the overlap of two MiSeq next-generation sequencing (NGS) datasets. **B)** Binding kinetics for DH1285 UCA (**B**) IgG to $Man_5GlcNAc_2$-enriched CH505.M5.G458Y Env trimer. **C-D)** Binding magnitude of DH1285 lineage members derived from NGS for CH505.M5.G458Y (blue), CH505.TF (red), or SARS-CoV2 NTD (gray). The UCA is composed of an inferred VH and VK from NGS of heavy and light chains. The NGS identified mutated DH1285 VHs are paired with the DH1285 light chain. DH1331 is a rhesus anti–SARS-CoV-2 receptor binding domain antibody. 2G12Rh is a chimeric antibody composed of a human light chain paired with a heavy chain made of human variable region fused to the gamma constant region of rhesus macaque. **E)** Multisequence alignment of DH1285 lineage members. Boxes show amino acids found in CH505 TF-reactive lineage members. Related to Figure 6.

### Table S1. CH235.12 Cryo-EM data collection and refinement statistics

| | 588AMS CH505.M5/293F | 603AMS CH505.M5/GnT1- |
|---|---|---|
| **Data Collection** | | |
| **Microscope** | FEI Titan Krios | FEI Titan Krios |
| **Voltage (kV)** | 300 | 300 |
| **Electron dose (e-/Å2)** | 65.94 | 66.71 |
| **Detector** | Gatan K3 | Gatan K3 |
| **Pixel Size (Å)** | 1.08 | 1.08 |
| **Defocus Range (μm)** | ~0.75-2.50 | ~0.75-2.50 |
| **Magnification** | 81000 | 81000 |
| **Micrographs collected** | 3189 | 3289 |
| **Reconstruction** | | |
| **Software** | cryoSparc | cryoSparc |
| **Particles** | 53398 | 50295 |
| **Symmetry** | C3 | C3 |
| **Box size (pix)** | 350 | 350 |
| **Resolution (Å) (FSC 0.143)*** | 3.7 | 4.1 |
| **Refinement (Phenix)** | | |
| **Protein residues** | 2988 | 3054 |
| **Chimera CC** | 0.72 | 0.66 |
| **R.m.s. deviations** | | |
| **Bond lengths (Å)** | 0.012 | 0.028 |
| **Bond angles (°)** | 2.038 | 2.110 |
| **Validation** | | |
| **Molprobity score** | 1.54 | 2.05 |
| **Clash score** | 0.38 | 3.72 |
| **Favored rotamers (%)** | 97.23 | 97.52 |
| **Ramachandran** | | |
| **Favored regions (%)** | 89.45 | 88.30 |
| **Allowed regions (%)** | 8.5 | 10.20 |
| **Disallowed regions (%)** | 2.05 | 1.50 |

*Resolutions are reported according to the FSC 0.143 gold-standard criterion

### Table S2. DH1285 Cryo-EM data collection and refinement statistics

| | DH1285 bound to CH505 Env Trimer Local Refinement | DH1285 bound to CH505 Env Trimer - 3 Fab bound state | DH1285 bound to CH505 Env Trimer - 2 Fab bound state |
|---|---|---|---|
| **PDB ID** | | | |
| **EMDB ID** | | EMD-27621 | EMD-27622 |
| **Data Collection and processing** | | | |
| **Microscope** | FEI Titan Krios | | |
| **Detector** | Gatan K3 | | |
| **Magnification** | 81000 | | |
| **Voltage (kV)** | 300 | | |
| **Electron exposure (e-/Å^2)** | 59.1 | | |
| **Defocus Range (µm)** | 2.4 to 0.8 | | |
| **Pixel size (Å)** | 1.08 | | |
| **Micrographs collected** | 7706 | | |
| **Reconstruction software** | cryoSPARC | | |
| **Symmetry imposed** | C1 | C3 | C1 |
| **Initial particle images (no.)** | 4,146,195 | | |
| **Final particle images (no.)** | 78,100 | 61,923 | 78,101 |
| **Map resolution (Å)** | 4.46 | 6.95 | 5.5 |
| **FSC threshold** | 0.143 | 0.143 | 0.143 |
| **Refinement** | | | |
| **Model resolution (Å)** | 5.1 | | |
| **FSC threshold** | 0.5 | | |
| **Model composition** | | | |
| **Nonhydrogen atoms** | 5016 | | |
| **Protein residues** | 603 | | |
| **R.M.S. deviations** | | | |
| **Bond lengths (Å)** | 0.005 | | |
| **Bond angles (°)** | 0.956 | | |
| **Validation** | | | |
| **MolProbity score** | 1.88 | | |
| **Clashscore** | 7.02 | | |
| **Favored rotamers (%)** | 98.31 | | |
| **Ramachandran plot** | | | |
| **Favored regions (%)** | 91.77 | | |
| **Disallowed regions (%)** | 0 | | |

*Resolutions are reported according to the FSC 0.143 gold-standard criterion
